# Genetic control of kinetochore-driven microtubule growth in *Drosophila* mitosis

**DOI:** 10.1101/2021.11.19.469214

**Authors:** Julia V. Popova, Gera A. Pavlova, Alyona V. Razuvaeva, Lyubov A. Yarinich, Evgeniya N. Andreyeva, Alina F. Anders, Yuliya A. Galimova, Fioranna Renda, Maria Patrizia Somma, Alexey V. Pindyurin, Maurizio Gatti

**Author notes:** Corresponding authors: Maurizio Gatti (MG) Alexey V. Pindyurin (AVP). These authors contributed equally to this work (Julia V. Popova, Gera A. Pavlova).

## Abstract

Centrosome-containing cells assemble their spindles exploiting three main classes of microtubules (MTs): MTs nucleated by the centrosomes, MTs generated near the chromosomes/kinetochores, and MTs nucleated within the spindle by the augmin-dependent pathway. Mammalian and *Drosophila* cells lacking the centrosomes generate MTs at kinetochores and eventually form functional bipolar spindles. However, the mechanisms underlying kinetochore-driven MT formation are poorly understood. One of the ways to elucidate these mechanisms is the analysis of spindle reassembly following MT depolymerization. Here, we used an RNA interference (RNAi)-based reverse genetics approach to dissect the process of kinetochore-driven MT regrowth (KDMTR) after colcemid-induced MT depolymerization. This MT depolymerization procedure allows a clear assessment of KDMTR, as colcemid disrupts centrosome-driven MT regrowth but allows KDMTR. We examined KDMTR in normal *Drosophila* S2 cells and in S2 cells subjected to RNAi against conserved genes involved in mitotic spindle assembly: *mast*/*orbit*/*chb* (*CLASP1*), *mei-38* (*TPX2*), *mars* (*HURP*), *dgt6* (*HAUS6*), *Eb1* (*MAPRE1/EB1*), *Patronin* (*CAMSAP2*), *asp* (*ASPM*) and *Klp10A* (*KIF2A*). RNAi-mediated depletion of Mast/Orbit, Mei-38, Mars, Dgt6 and Eb1 caused a significant delay in KDMTR, while loss of Patronin had a milder negative effect on this process. In contrast, Asp or Klp10A deficiency increased the rate of KDMTR. These results coupled with the analysis of GFP-tagged proteins (Mast/Orbit, Mei-38, Mars, Eb1, Patronin and Asp) localization during KDMTR suggested a model for kinetochore-dependent spindle reassembly. We propose that kinetochores capture the plus ends of MTs nucleated in their vicinity and that these MTs elongate at kinetochores through the action of Mast/Orbit. The Asp protein binds the MT minus ends since the beginning of KDMTR, preventing excessive and disorganized MT regrowth. Mei-38, Mars, Dgt6, Eb1 and Patronin positively regulate polymerization, bundling and stabilization of regrowing MTs until a bipolar spindle is reformed.

**Author summary:** The mitotic spindle is a microtubule (MT)-based molecular machine that mediates precise chromosome segregation during cell division. Both *Drosophila* and human cells assemble their spindles exploiting two main classes of MTs: MTs nucleated by the centrosomes (MT nucleating organelles) and MTs generated at or near the kinetochores (the chromosome-associated structures that bind the spindle MTs). Cells of both species can assemble a functional mitotic spindle in the complete absence of centrosomes, but the mechanisms underlying this process are still poorly understood. We used *Drosophila* S2 cells as model system to analyze spindle reassembly following colcemid-induced MT depolymerization. MT regrowth (MTR) after colcemid treatment was particularly informative as this drug disrupts the MT nucleating ability of the centrosomes but allows kinetochore-driven MTR (KDMTR). We analyzed KDMTR in normal cells and in cells subjected to RNA interference (RNAi)-mediated depletion of 8 different evolutionarily conserved proteins involved in spindle assembly, and identified proteins that either promote or delay KDMTR. These results coupled with the analysis of proteins localization during spindle reassembly allowed us to integrate the current model on the role of kinetochore-driven MT growth in spindle formation.

## Introduction

The spindle is a microtubule (MT)-based highly dynamic molecular machine that mediates precise chromosome segregation during both mitosis and meiosis. To form a spindle, centrosome-containing cells generate MTs in three main cellular locations: at the centrosomes, near chromosomes and/or at kinetochores, and within the spindle through the augmin-mediated pathway (reviewed in [1-4]). MTs are always nucleated by the γ-tubulin ring complexes (γ-TuRCs), which are embedded in the centrosomes, enriched in the vicinity of the kinetochores or associated with the walls of the spindle MTs by interaction with augmin (reviewed in [3, 5]). Studies carried out in mammalian tissue culture cells and in different types of *Drosophila* somatic cells have shown that chromosome/kinetochore-driven MT formation is sufficient for the assembly of a functional spindle, but to date little is known about the factors that govern the growth of kinetochore-dependent MTs (reviewed in [3, 4, 6, 7]).

Early studies using *Xenopus* oocyte extracts revealed that chromatin has the ability of driving MT growth and bipolar spindle formation (reviewed in [8]). In addition, mammalian cells are able to form bipolar spindles after centrosome ablation with laser microsurgery [9]. Consistent with these results, *Drosophila* mutants devoid of centrosomes, or with centrosomes with strongly reduced MT nucleating activity [e.g. *asl* (*CEP152*), *Sas-4* (*CENPJ*), *cnn* (*CDK5RAP2*) and *spd-2* (*CEP192*) mutants; unless mentioned otherwise, here and henceforth the human ortholog of the fly gene or protein is reported within brackets], can assemble functional mitotic spindles and develop to adulthood [6, 10-14]. Centrosomal MTs are also dispensable for spindle formation in *Drosophila* tissue culture cells. For example, S2 cells subjected to RNAi-mediated depletion of centrosomal components, such as Cnn, Sas-4 and Spd-2, assemble functional anastral spindles [15-17]. Thus, centrosomes and astral MTs appear to be dispensable for the assembly of a functional spindle in both mammalian and *Drosophila* somatic cells.

Three main approaches are currently used to analyze chromatin/kinetochore-driven MT formation. A first approach involves direct examination of kinetochore fibers (k-fibers) formation from unattached kinetochores in live centrosome-containing cells expressing GFP-tubulin (reviewed in [1]). A second approach exploits systems devoid of functional centrosomes such as *Xenopus laevis* extracts or cells deficient of critical centrosomal proteins required for MT nucleation. A third approach consists in the analysis of spindle MTs regrowth (MTR) after cold-or drug-induced MT depolymerization. In *Xenopus laevis* extracts, chromatin or DNA-coated beads stimulate MT nucleation and polymerization along their entire surface (reviewed in [8]). Similarly, in *Drosophila* embryos MTR occurs throughout mitotic chromatin [18]. In contrast, in S2 cells and human cells MT growth is restricted to the kinetochore regions [19-25] (and this study).

Studies on mitosis in *Xenopus* extracts and vertebrate cells have shown that the GTP-bound form of Ran GTPase (RanGTP) stimulates chromatin-induced MTs growth. RanGTP is generated in the vicinity of chromosomes by RCC1, a chromosome-associated RanGTP exchange factor (reviewed in [8]). In both *Xenopus* and mammalian systems, RanGTP forms a gradient highly concentrated around the chromosomes that positively regulates several MT-associated proteins including Aurora A, TPX2, HURP, Aurora B, INCENP and Nup107-160 (reviewed in [26]). The role of RanGTP in chromosome-driven MT growth has been also studied in *Drosophila* embryos and different types of fly somatic cells. Although S2 tissue culture cells form a RanGTP gradient around the chromosomes, RNAi-mediated depletion of >95% of RCC1 does not affect spindle assembly and functioning, and, consistently, it does not result in defective kinetochore-driven MT growth [27]. However, *Drosophila* embryos injected with a dominant negative form of Ran are severely defective in chromosome-driven MTR after cold-induced depolymerization [18, 28]. Thus, it is currently unclear whether chromosome-associated MT polymerization in S2 cells requires a minimal concentration of RanGTP, or whether it is RanGTP-independent.

The current model on the role of kinetochores in spindle assembly is largely based on the analysis of mitosis in centrosome-containing *Drosophila* S2 cells expressing GFP-tagged tubulin. Careful observations on mitosis in these live cells, accompanied by laser microsurgery experiments, suggested that the plus ends of short chromatin-induced MTs are captured by the kinetochores and continue to polymerize there, leading to growing bundles of MTs with the minus ends that are pushed away from the kinetochores [20]. These growing bundles, which will give rise to the k-fibers, interact with the astral MTs and eventually coalesce to form a bipolar spindle [20, 29]. A similar model applies to both *Drosophila* and human cells that form bipolar spindles in the absence of centrosomes or reassemble a spindle after MT depolymerization. However, in these cases the kinetochore-driven k-fibers coalesce at the spindle poles through a centrosome-independent mechanism that exploits MT minus end-directed motors and minus end binding proteins [7, 29].

Studies on human cells, *Xenopus-*derived systems and *Drosophila* identified several proteins that control chromatin/kinetochore-driven MTs growth. For example, this process is inhibited by depletion of TPX2, HURP, Aurora A, Aurora B or INCENP, whose deficiency does not impair MT nucleation from the centrosomes [21, 22, 30-32]. In *Drosophila* embryos, chromosome-associated MTR is prevented by depletion of the HURP homologue Mars, but not by the TPX2 homologue Mei-38 [18]. In *Drosophila* larval brain cells, kinetochore-driven MTR (KDMTR) is inhibited by loss of Misato (Mst), a protein that interacts with the Tubulin Chaperone Protein-1 (TCP-1) complex and the Tubulin Prefoldin complex, which are also required for KDMTR [33, 34]. Other factors required for efficient KDMTR in *Drosophila* somatic cells are Ensconsin (Ens), an MT-binding protein homologous to the human MAP7 [35], γ-tubulin and the Msps (TOGp) MT polymerase [23]. Finally, in mammalian cells KDMTR is hampered by loss of the MT minus end binding MCRS1-KANSL1-KANSL3 complex [36, 37] and by failure of Nup107-160-dependent tubulin recruitment at kinetochores [38].

Another conserved factor that promotes chromatin-induced MT formation is augmin. Augmin is an 8-protein complex that binds the lateral walls of spindle MTs and recruits γ-TuRCs that nucleate additional MTs, “augmenting” the spindle MT density [39-41]. In human cells, *Drosophila* S2 cells and *Drosophila* embryos, augmin is required for chromosome-driven MT formation and efficient assembly of k-fibers [18, 23, 39, 42-45]. Interestingly, *Drosophila* mutants in the *wac* and *msd1* genes, each of which encodes an augmin subunit, are viable and do not exhibit defective spindles in larval brains. However, flies homozygous for both *cnn* and *msd1* mutations are lethal and display highly aberrant spindles [46]. These results suggest that in *msd1* mutants there is a reduced chromosome-driven MT generation, which is sufficient for spindle assembly in the presence of MTs nucleated by the centrosomes, but insufficient in the absence of centrosomal MTs.

All these studies indicate that the mechanisms of KDMTR can be dissected using a genetic approach. Here, we use *Drosophila* S2 cells to analyze MTR after colcemid treatment. We examine this process in normal cells and in cells subjected to RNAi-mediated depletion of different evolutionarily conserved proteins. Specifically, we analyze MTR in cells depleted of (i) proteins that have been already implicated in KDMTR such as Mars (HURP), Mei-38 (TPX2) and the augmin subunit Dgt6 [18, 21-23, 31, 32], (ii) the plus end-associated factors Eb1 (MAPRE1/EB1) and Mast (CLASP1) (reviewed in [47]) (*mast/orbit*, whose official FlyBase name is *chromosome bows, chb*, will be henceforth designated as *mast*), (iii) the minus end binding Patronin (CAMSAP2) and Asp (ASPM) (reviewed in [48]) and (iv) the Klp10A kinesin-like protein (KIF2A) that depolymerizes the MT minus ends (reviewed in [49]). We show that depletion of some these proteins (Mast, Mars, Mei-38, Dgt6 and Eb1) down-regulates KDMTR. In contrast, loss of Asp or Klp10A leads to an increased mass of regrowing MTs compared to control. We also examined the localization of GFP-tagged Mast, Mars, Mei-38, Eb1, Patronin and Asp during MTR. This analysis revealed that these proteins exhibit different localizations during KDMTR, which are likely to reflect their specific roles in the process. Collectively, our results define the modes of spindle reassembly after colcemid-induced MT depolymerization and identify proteins that either enhance or reduce KDMTR.

## Results

To investigate the mechanisms of KDMTR, we used a reverse genetic approach. We performed RNAi in *Drosophila* S2 cells against 8 genes encoding proteins required for mitotic spindle assembly and then examined RNAi cells for spindle reformation after MT depolymerization with colcemid. Specifically, we focused on *mast, mars, mei-38, Eb1, dgt6, Patronin, asp* and *Klp10A*. Before performing the MTR experiments, we checked the efficiency of RNAi by quantifying the reduction of target mRNA level and examining the phenotypic consequences thereof.

### Efficiency of RNAi

We determined the level of target mRNA after 5 days incubation with the corresponding double-stranded RNA (dsRNA). In all cases (except *Patronin*) we found mRNA levels ranging from 3 to 8 % of the untreated cells level; *Patronin* mRNA was reduced to 28% of control (S1 Table). We then examined the effect of RNAi in fixed cells immunostained for α-tubulin and the centrosome marker Spd-2 and counterstained for DNA with DAPI. Consistent with previous results [50-53], we found that RNAi-mediated depletion of the Klp10A MT depolymerase or of the Asp minus end binding protein results in longer late prometaphase/metaphase spindles compared to those observed in control cells (S1 Figure). Klp10A accumulates at the kinetochores, the centrosomes and the spindle poles but it thought to act mainly at the poles [54, 55]. *Klp10A* RNAi cells showed frequent monopolar spindles and a modest but significant increase in the frequency of prometaphase-like cells with elongated spindles (PMLES) (S1 Table; see also [56]). PMLES (or pseudo ana-telophases, PATs) have been previously observed in cells where the kinetochore-MT interaction was compromised, such as those depleted of the centromeric histone Cid (CENPA) or the Ndc80 kinetochore protein [16, 57]. Asp binds the MT minus ends [58] and is enriched at both the spindle poles and the extremities of the central spindle [51, 53, 59, 60]. In agreement with these studies, we found that Asp-depleted cells exhibit broad spindle poles that often show centrosome detachment, and defects in chromosome alignment and segregation (i.e. PMLES; S1 Table).

Cells depleted of Mast, Mars, Mei-38, Dgt6, Eb1 or Patronin are all characterized by spindles significantly shorter than those of control cells (S1 Figure and [23]) but exhibit very different mitotic phenotypes. The most dramatic phenotype was observed in cells depleted of Mast, which mediates the incorporation of tubulin dimers into the plus ends of the MTs embedded in the kinetochores [61]. As described previously [62, 63], in *mast* RNAi cells most spindles collapse forming monopolar figures, while the short bipolar spindles show defective chromosome alignment and segregation (S1 Figure, S1 Table). Eb1 is enriched at the plus ends of growing MTs and its depletion results in short and malformed spindle and defective chromosome segregation [64]. In line with these results, we found that *Eb1* RNAi cells exhibit many morphologically abnormal spindles and a higher frequency of PMLES compared to control (S1 Figure, S1 Table). Frequent PMLES and monopolar spindles were also observed in cells depleted of the augmin component Dgt6, as described previously [23]. Depletion of Patronin, which binds the MT minus ends, resulted in short spindles often associated with multiple centrosomes (S1 Figure, S1 Table; see also [53]). The spindles of both *mei-38* and *mars* RNAi cells were shorter than control spindles but did not show gross morphological defects, in agreement with previous results [65-67]. However, *mei-38* RNAi cells, displayed an increase in monopolar spindles compared to control. A small but significant increase in monopolar spindles was also observed in *mars* RNAi cells, together with mild defects in chromosome segregation (S1 Figure, S1 Table).

### KDMTR after colcemid-induced depolymerization

To address the roles of the genes of interest in KDMTR, we analyzed the effects of their deficiency following colcemid-induced MT depolymerization. Previous work has shown that after cold-induced MT depolymerization the centrosomes of S2 cell prometaphases and metaphases (henceforth PRO-METs) are normally placed at the presumed location of the depolymerized spindle poles, where they nucleate MT asters [23, 68]. In contrast, after colcemid treatment, the centrosomes are no longer able to drive astral MT regrowth and in most cells were freely floating in the cytoplasm ([68]; see also Figures 1 and 2). As a result, in many cases one or both poles of the reformed spindles were not associated with the centrosomes [68]. Thus, by analyzing MTR after colcemid-treatment we are in fact studying KDMTR in the absence of centrosomal activity.

**Figure 1.**
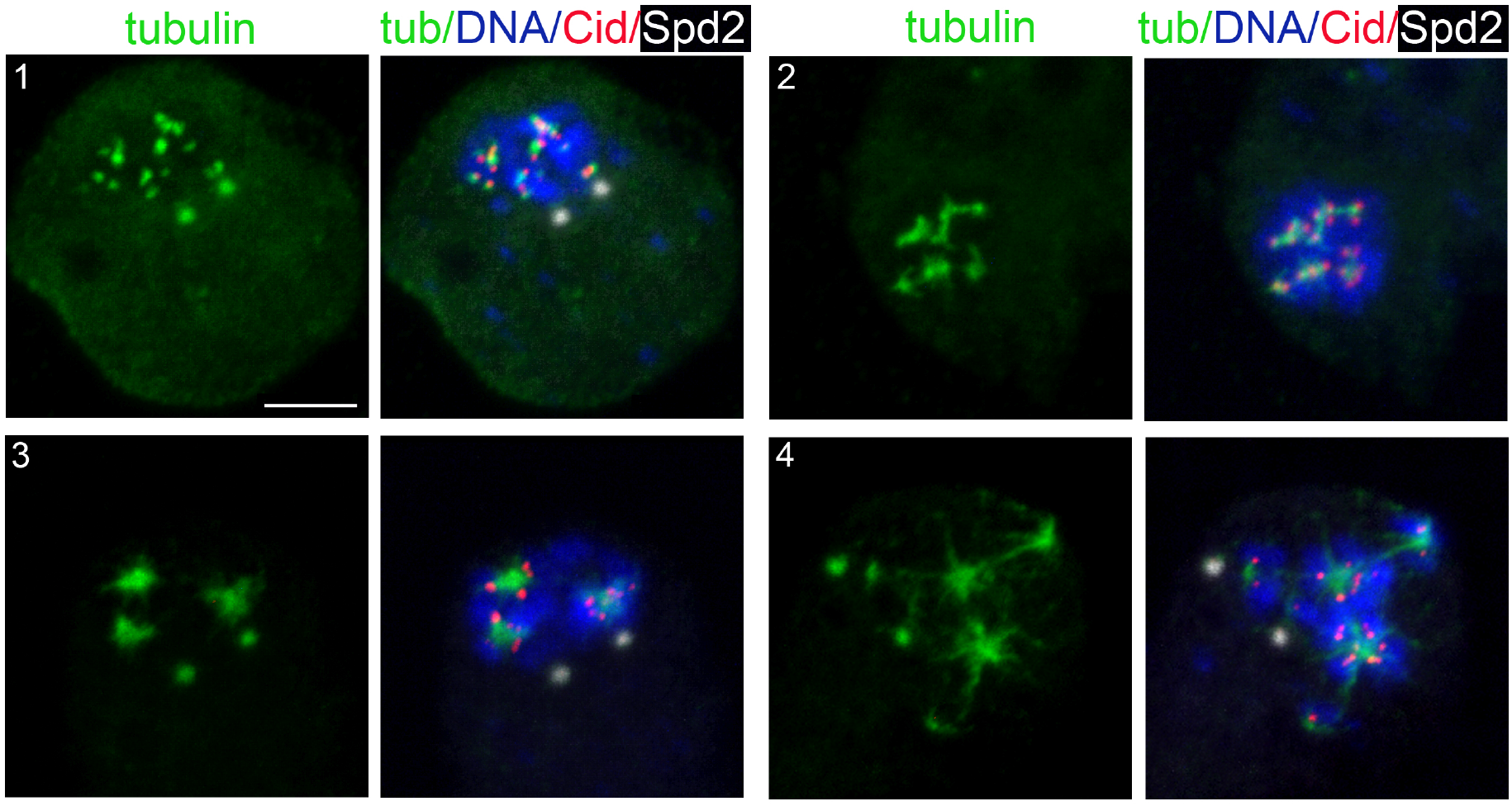
Representative examples of KDMTR after colcemid-induced MT depolymerization. S2 cells were fixed and stained with anti-Cid (red), anti-α-tubulin (green) and anti Spd-2 (white) antibodies and counterstained for DNA (DAPI, blue). Panel 1 shows an initial MTs regrowth phase with small tubulin signals associated with Cid signals (centromeres). In Panel 2, some of the initial signals have elongated and appear as MT bundles. Panels 3 and 4 illustrate a further regrowth phase, with MTs forming clusters/asters, which are surrounded by centromeres. Scale bar, 5 µm.

**Figure 2.**
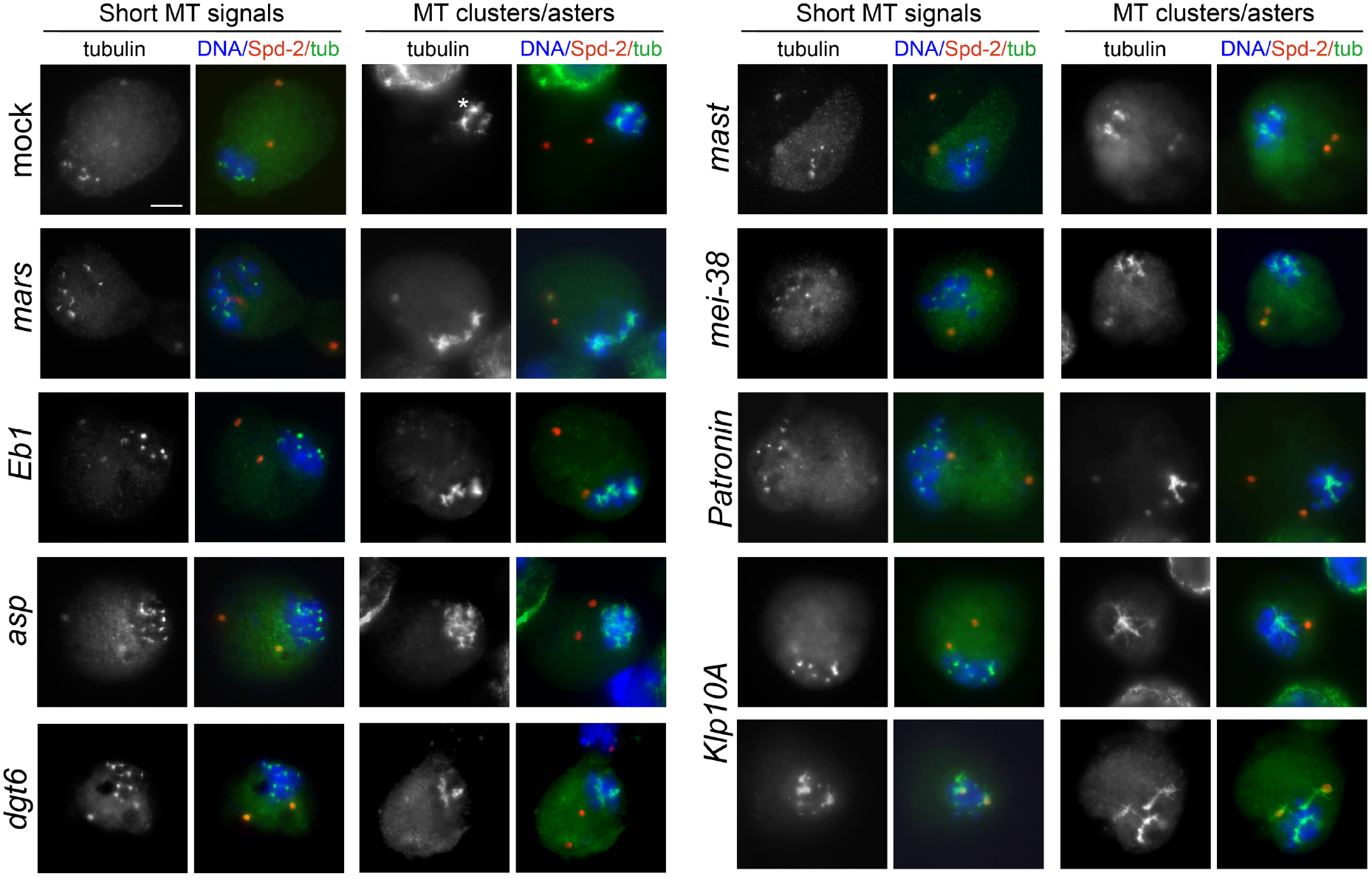
Examples of representative MTR figures in mock-treated and RNAi cells following colcemid-induced (3 h, 4.5 µg/ml) MT depolymerization. S2 cells were fixed 30 min after colcemid washout and stained for α-tubulin (green), centrosomes (Spd-2, red) and DNA (DAPI, blue). The cells show several combinations of MTR figures: (i) small and double roundish signals emanated by one or two kinetochore, respectively (see Figure 1), (ii) relatively long MT bundles; (iii) MT cluster/asters. Note that after colcemid treatment the prometaphase and metaphase chromosomes of both control and RNAi cells remain clustered, while in most cases the centrosomes are not associated with regrowing MTs and appear to float freely in the cytoplasm. In contrast, in a substantial fraction of *Klp10A* RNAi cells the centrosome are surrounded by MTs. Control and RNAi cells show morphologically comparable MTR figures but differ in the frequency of these figures, as well as in the fluorescence intensity of regrowing MTs (see Table 2). Scale bar, 5µm.

Control and RNAi (*mast, mars, mei-38, Eb1, Patronin, dgt6, asp* and *Klp10A*) cells were treated for 3 h with colcemid. Cells were then accurately washed to remove colcemid and fixed after 20, 30, 45 and 75 min after removal of the drug. Some cells were fixed after 3 hours of colcemid treatment without removal of the drug, to check the degree of MT depolymerization (time 0). All cells were immunostained for both α-tubulin and the centrosomal marker Spd-2 and counterstained with DAPI. We limited our observations to PRO-METs, as the kinetochores of cells in these mitotic phases have the ability to drive k-fiber formation and bipolar spindle reassembly [68].

At time 0, in more than 95% of the PRO-METs, the spindle was completely depolymerized, indicating that none of the RNAi treatments affects colcemid-induced MT depolymerization. At 20, 30 and 45 min after colcemid removal in *mast, mars, mei-38, Eb1* and *Patronin* RNAi cells the frequencies of PRO-METs showing KDMTR were significantly lower than in control (Table 1). *dgt6* RNAi cells with KDMTR were less frequent than in control only at the 20 and 30 min fixation times (Table 1). In *asp* and *Klp10A* RNAi cells the frequencies of PRO-METs showing KDMTR were broadly similar to the control frequencies with two exceptions. At 20 min, the frequency of *Klp10A* RNAi cells showing KDMTR was higher than in control, while at 30 min the frequency of Asp-depleted cells with KDMTR was slightly but significantly lower than in control. After 75 min of MT regrowth, nearly all RNAi and control cells displayed KDMTR (Table 1). These results suggest that cells depleted of Mast, Mars, Mei-38, Eb1, Patronin or Dgt6 are defective in KDMTR. However, we would like to note that the frequency of cells showing KDMTR reflects the capability of mitotic cells to initiate but not to sustain and promote this process during spindle reassembly.

**Table 1.**
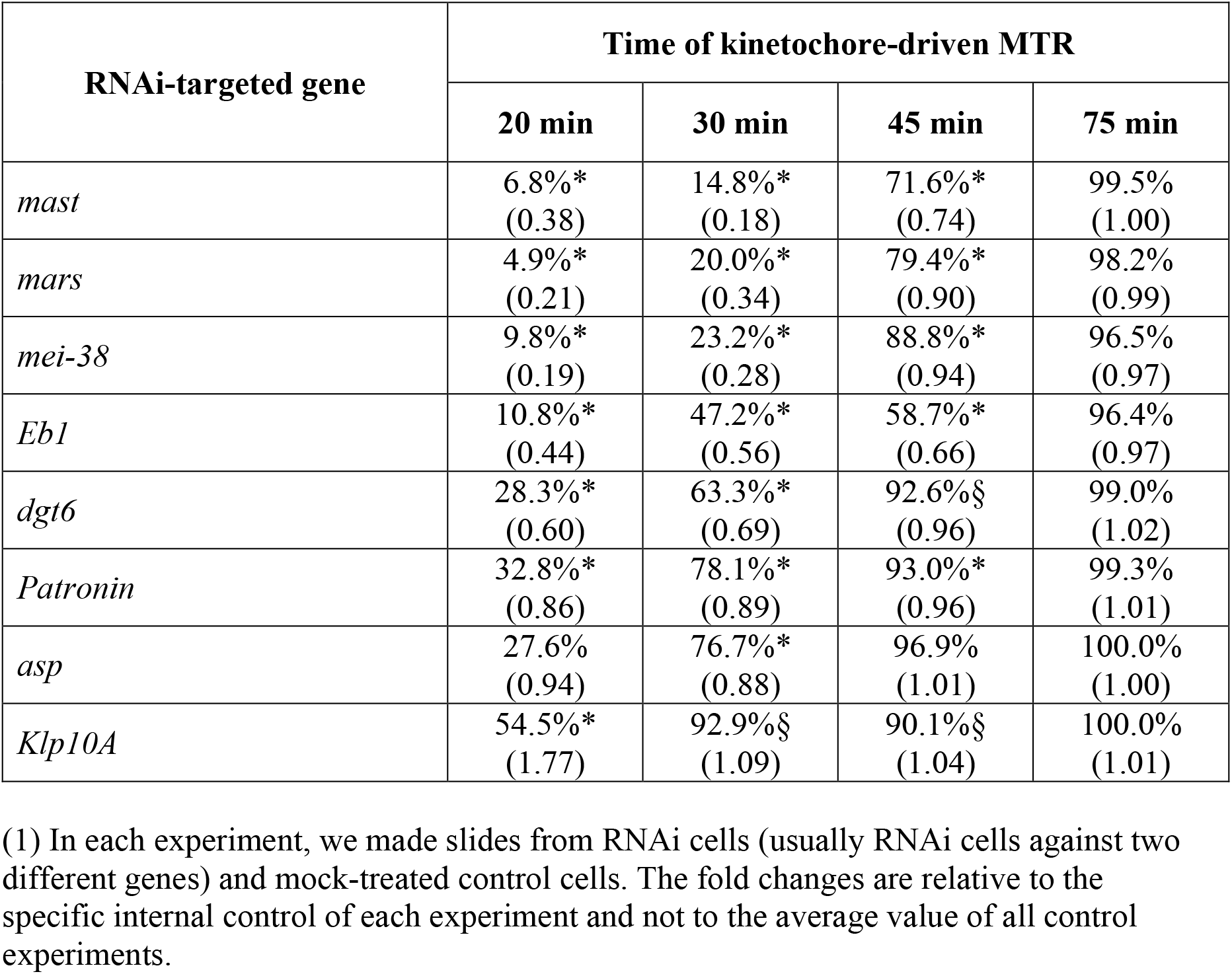
Mean frequencies and fold changes (between brackets) relative to control (1) of cells showing KDMTR at different times after colcemid washout. In all cases, except *Klp10A*, data are from at least 3 independent experiments; for *Klp10A* RNAi cells data are from 2 independent experiments. § and *, significant in χ^2^ test with p ≤ 0.05 and ≤ 0.01, respectively.

To gather additional information about KDMTR, we analyzed the pattern of MTR. After colcemid-induced MT depolymerization, MT repolymerization in control cells started at kinetochores, which became tightly associated with small dot-like tubulin signals (Figure 1). These tubulin dots then expanded, forming tubulin bundles and tubulin aggregates that often appeared as aster-like structures. However, these structures did not contain centromeres at their centers but were instead surrounded by centromere signals (Figures 1 and 2). In some cells, centrosomes were associated with weak tubulin signals (Figures 1 and 2), but never showed astral MT regrowth as occurs after cold-induced MT depolymerization [23, 68]. We distinguished three types of short KDMTR signals: kinetochore-associated tubulin dots and double dots (or rods of the size of a double dot) corresponding to the initial MTR from a single kinetochore or both sister kinetochores (Figures 1 and 2). Other small signals were relatively long MT bundles resulting from the elongation of the “double dots” (Figures 1 and 2). With progression of MTR, these small signals expanded forming tubulin clusters that often appeared as aster-like structures (Figures 1 and 2). These structures then coalesced and emanated long MT bundles, resulting in different types of MT intermediate arrangements, including many umbrella-like formations in which the regrowing bundles converge on one side and diverge on the other. Most likely, each of these intermediate figures will give rise to a bipolar anastral spindle. For example, as documented in previous studies (see for example [15, 69]) the divergent MT bundles of the umbrella-like structures elongate and progressively merge to form a second spindle pole. Reformation of a bipolar spindle after colcemid-induced MT depolymerization is shown in S2 Figure; the structure of the MT clusters/asters is described below together with Asp-GFP and Mast-GFP localization during spindle reassembly.

At 20 min fixation time the control cells with KDMTR are relatively few and frequently show very small tubulin signals. In contrast, at 45 min fixation time tubulin signals are large and abundant and often exhibit complex morphologies. Thus, to quantitate KDMTR in control and RNAi cells we focused on PRO-METs fixed 30 min after colcemid removal, as these cells exhibit clear kinetochore-associated tubulin signals that can be examined for both morphology and fluorescence intensity. Importantly, we found that control and RNAi PRO-METs exhibit very similar KDMTR figures, which, however, vary in frequency according to the RNAi treatment. We distinguished three types of MTR figures: (i) very short kinetochore-associated MT bundles (the single and “double spot” signals described above), (ii) relatively long MT bundles, and (iii) MT clusters/asters. In *mast, mars, mei-38, dgt6* and *Eb1* RNAi cells, the frequencies per cell of elongated MT bundles and MT clusters/asters, which represent advanced stages of KDMTR, were significantly reduced compared to controls (Table 2; S3 Figure). These RNAi cells also showed a concomitant significant reduction of chromosome-associated tubulin fluorescence (CATF) (Table 2; S4 Figure). In *Patronin*-depleted cells, the frequencies of MT bundles and clusters/asters were slightly but not significantly reduced compared to control. However, in *Patronin* RNAi cells CATF was significantly lower than in control (Table 2; S3 and S4 Figures). In contrast, Asp- and Klp10A-depleted cells showed significant increases in both MT clusters/asters and CATF compared to controls (Table 2; S3 and S4 Figures). The effect of Asp depletion on KDMTR was completely unexpected and suggests a role for this protein in the process of kinetochore-driven MT growth in unperturbed cells.

**Table 2.**
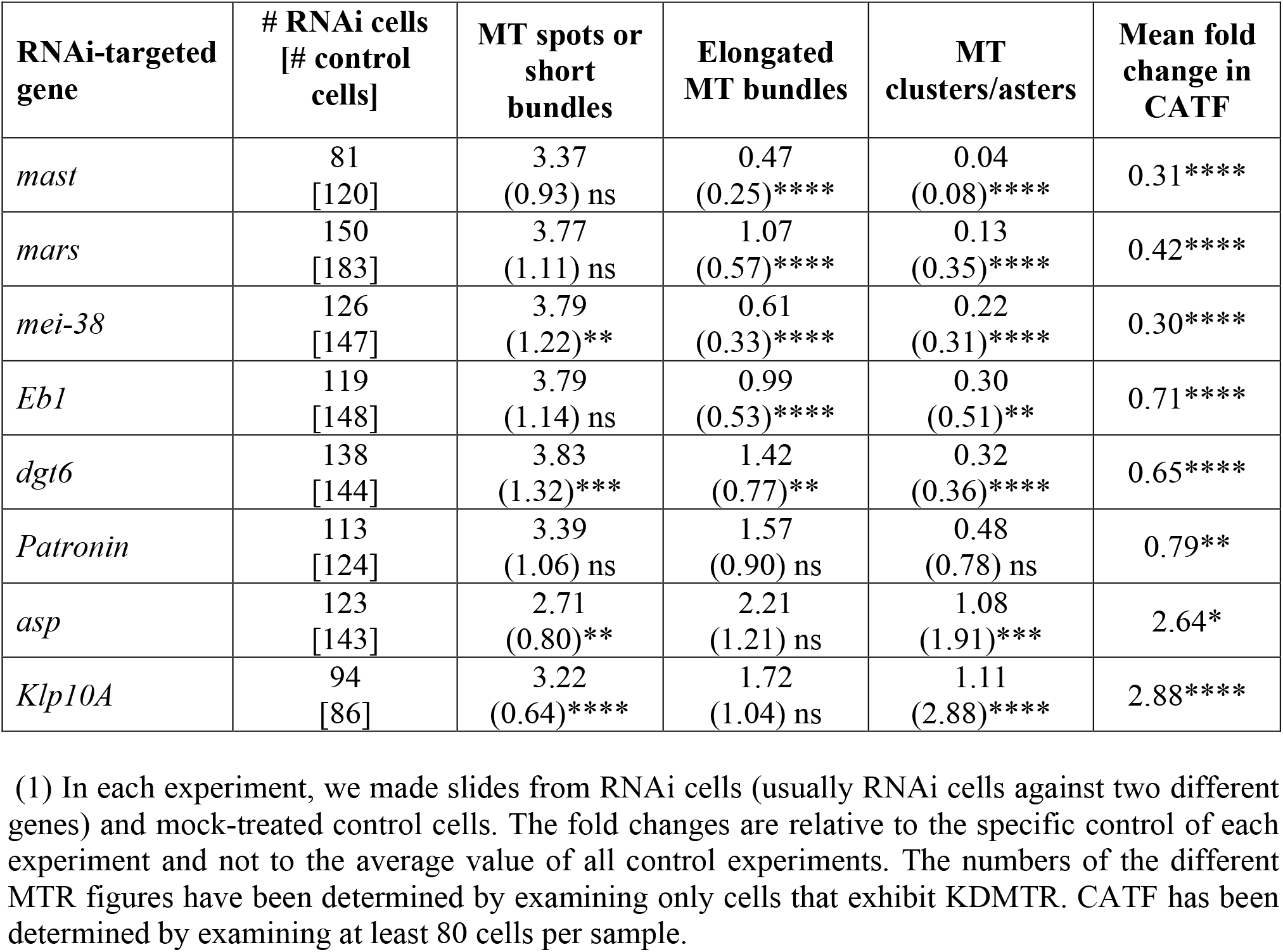
Mean numbers of the indicated MTR figures per cell observed 30 min after colcemid washout. Between round brackets are the mean fold changes of these numbers relative to control (1). The mean fold changes of chromosome-associated tubulin fluorescence (CATF) relative to control (1) are also indicated. In all cases, except *Klp10A*, data are from at least 3 independent experiments; for *Klp10A* RNAi cells data are from 2 independent experiments. The numbers of RNAi and control cells examined for definition of the MTR pattern are indicated in the first column. Data have been analyzed with the Mann-Whitney *U* test; ns, not significant; *, **, ***, **** significant with p ≤ 0.05, 0.01, 0.001, and 0.0001, respectively.

Interestingly, in Klp10A-deficient cells a large fraction (55%) of the centrosomes were associated with strong tubulin signals (Figure 2). This event was seldom observed in control cells and in the other RNAi cells, which showed only very weak tubulin signals associated with the centrosomes (Figure 2). This observation provides insight into why colcemid-treated centrosomes lose their ability to drive MT growth, while retaining normal levels of pericentriolar material (PCM) proteins such as Spd-2. It is possible that these centrosomes retain their MT nucleating ability and that the regrowing MTs are particularly sensitive to the Klp10A MT depolymerase. Alternatively, colcemid-treated centrosomes might have lost the ability to shield the minus ends of the MTs they nucleate from the activity of Klp10A. Whatever the explanation, our findings underlie an interesting aspect of centrosome biology, which deserves future studies.

### Localization of selected spindle-associated proteins during KDMTR

To investigate the roles of the proteins that regulate KDMTR, we analyzed their localization during the process of spindle regrowth after colcemid treatment. In a previous study on MTR after cold exposure, we showed that the bundles of reforming MTs are uniformly enriched in Dgt6 since the beginning of their formation [23]. Here, we analyzed the localization of Mast, Mars, Mei-38, Eb1, Patronin and Asp. We generated S2 cell lines stably expressing Cherry-tubulin and each the protein of interest marked with GFP, both under the control of a copper-inducible promoter. The lines expressing the GFP fusions of the Patronin and Asp were described in a previous study [53]. The other lines were generated during the present investigation and their precise features are reported in Materials and Methods.

After induction of the transgenes by copper sulphate, Cherry-tubulin and the GFP-tagged proteins were visualized either in living cells or in fixed cells stained with anti-GFP and anti-α-tubulin antibodies and counterstained with DAPI. We found a perfect correspondence between the staining pattern of live and fixed cells, although in some cells tubulin staining was brighter after fixation and immunostaining compared to living cells. In all cases, the localization of GFP-tagged proteins in untreated cells was fully consistent with that observed in previously reports. Mast-GFP was bound to all spindle MTs and specifically enriched at the kinetochores, the spindle poles and the telophase spindle midzone (S5 Figure; see also [70]). Mei-38-GFP was uniformly distributed on the spindle MTs during all mitotic phases (S6 Figure; see also [67]). Mars-GFP too was uniformly distributed on PRO-MET spindles, but it was absent from the telophase central spindle (S7 Figure; see also [66, 71]). Eb1-GFP was associated with all spindle MTs and enriched at the growing MT plus ends (S8 Figure; see also [64]). Asp-GFP localized to all spindle MTs and concentrated at the spindle poles and the extremities of the telophase central spindle that are enriched in MT minus ends (S9 Figure; see also [51, 59, 72]). For Patronin-GFP, we confirmed the peculiar behavior we described previously [53]. In several live prometaphases, Patronin-GFP was associated with the entire spindle that appeared as a weakly and uniformly stained structure. However, these prometaphases suddenly showed brightly fluorescent Patronin-GFP signals associated with short MT bundles located near the chromosomes. These bright signals extended towards the cell poles along preexisting MT bundles (probably k-fibers) and stopped growing just before reaching the poles. Consistent with this finding, fixed prometaphases, displayed both GFP-stained and unstained MT bundles [53]. This behavior cannot be a consequence of Patronin-GFP overexpression as both dully fluorescent PRO-METs and those containing the bright MT bundles are likely to contain the same amounts of Patronin-GFP. We hypothesize that when GFP-tagged Patronin binds the k-fibers and move towards the spindle poles, it changes conformation exposing its GFP moiety. This conformational change would light up the k-fibers and would also result in a strong reaction with the anti-GFP antibody.

To gather information on the specific roles of the proteins that regulate KDMTR, we examined their behavior during MTR after colcemid-induced depolymerization. The cell lines carrying Cherry-tubulin and the GFP-tagged protein of interest, both under the control of a copper-inducible promoter, were treated for 16-22 h with copper sulfate, exposed to colcemid for 3 h, washed and then fixed at 30, 45, 90 and 120 min after colcemid removal. Some cells were fixed at the end of colcemid treatment before removal of the drug (time 0) The multiple fixation times permitted us to analyze different stages of MTR, ranging from the very early stages consisting of very small tubulin dots to the late stages containing long MT bundles often converging into a completely reformed bipolar spindle.

Following this experimental design, we first examined the localization of the minus end binding Asp protein. Surprisingly, at time 0, Asp-GFP was concentrated on the kinetochores of 50% of the PRO-METs (n = 200; Figure 3). Notably, in normal cells, Asp-GFP never accumulates on the kinetochores and it is always enriched at the spindle poles (S9 Figure). Thus, our observations suggest that upon MT depolymerization Asp redistributes in the cytoplasm and interacts with the kinetochores. At the very early MTR stages Asp-GFP was still accumulated at the kinetochores, showing partial co-localization with the initial tubulin foci. Asp-GFP was then found at the center of the tubulin cluster/asters (Figure 4). With progression of MTR, Asp-GFP associated with the extended regrowing MT bundles and accumulated at their minus ends (Figure 4A-4; see also S10 Figure). This finding is consistent with previous work showing that in normal spindles Asp moves along the MTs towards the spindle poles [60, 73]). Importantly, when the Asp-GFP signal was located at the center of the tubulin clusters/asters, the Cid (CENPA) centromeric signals were not at the center of these structures but were instead surrounding them (Figure 4B). These results suggest that Asp localizes at the MT minus ends at all regrowth stages, and that the aster-like structures are formed by the coalescence of the Asp-associated minus ends of the MTs emanating from kinetochores. Thus, these MT clusters/asters have features in common with the spindle poles and can be regarded as mini spindle poles.

**Figure 3.**
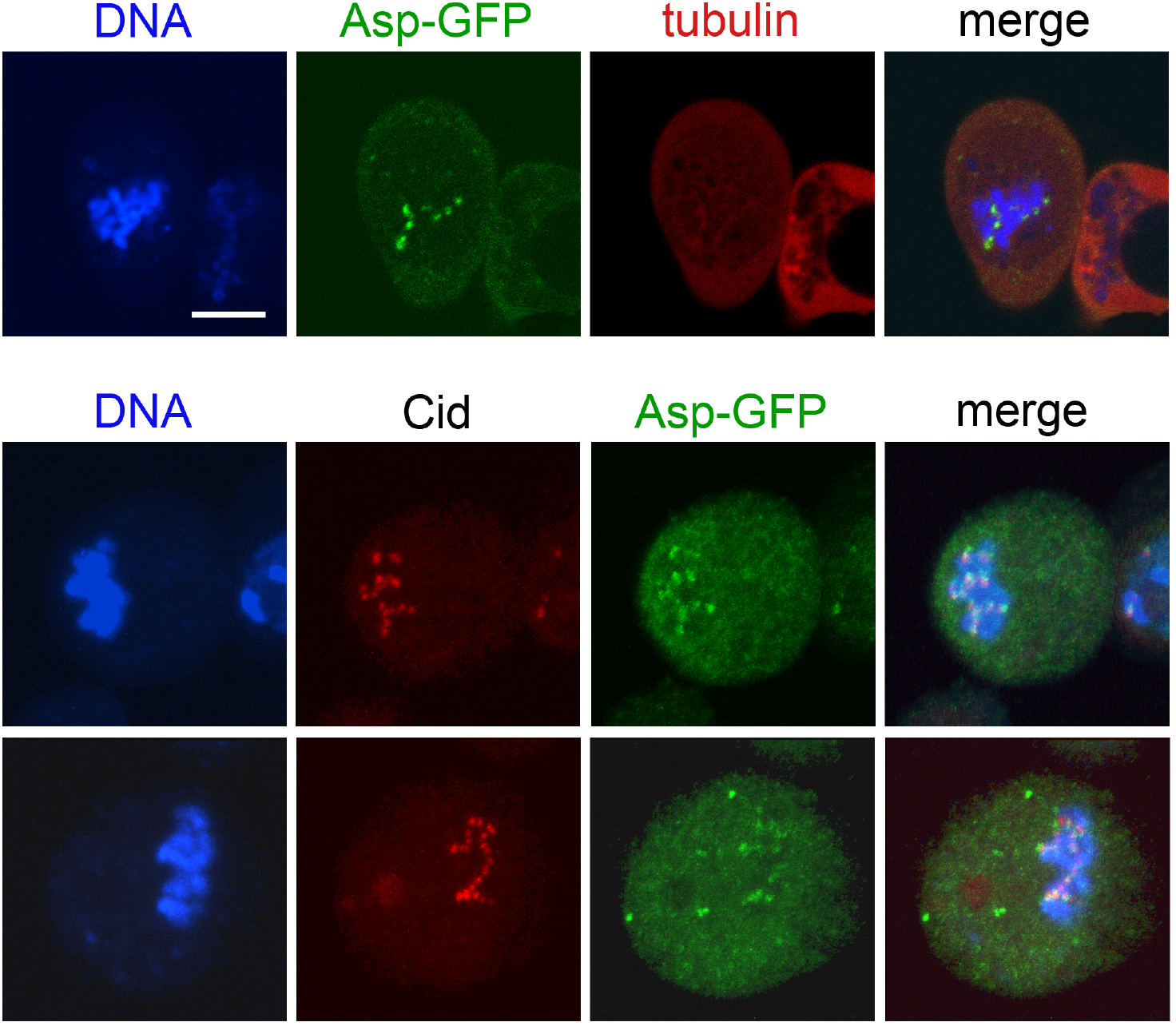
Asp-GFP localization in prometaphases exposed to colcemid for 3 h and photographed before colcemid washout. Top panels, a live S2 cell expressing Asp-GFP (green) and Cherry-tubulin (red) stained with the vital DNA stain Hoechst 33342 (blue). Bottom panels, representative fixed S2 cells expressing Asp-GFP stained with anti-GFP antibodies (green), anti-Cid antibodies (red) and DNA (DAPI; blue). Note that Asp-GFP is strictly associated with kinetochores. Scale bar, 5 µm

**Figure 4.**
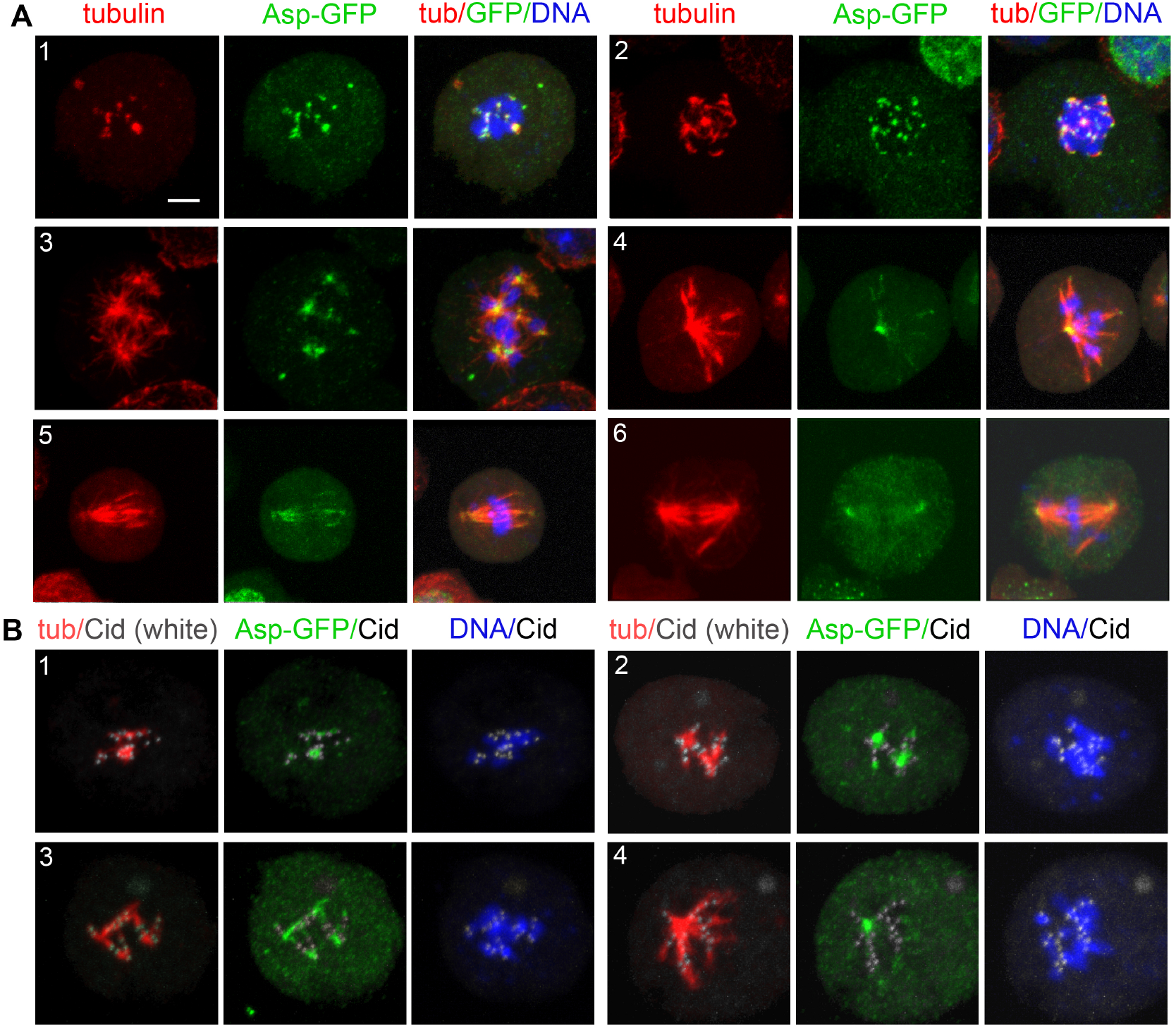
Asp-GFP localization during KDMTR after colcemid-induced MT depolymerization. **A**. S2 cells expressing both Asp-GFP and Cherry-tubulin were fixed and stained with anti-α-tubulin (red) and anti-GFP (green) antibodies and counterstained for DNA (DAPI, blue). Panel A1 shows an early KDMTR stage; panels A2 and A3 show more advanced KDMTR phases characterized by the presence of MT clusters/asters. Note that Asp-GFP is localized at the center of these MT structures, suggesting that they correspond to mini spindle poles. Panels A4-6 illustrate one of the modalities of spindle reformation after colcemid-induced MTs depolymerization. Panel A4 shows a representative monopolar spindle with Asp-GFP concentrated at the pole. In this regrowing spindle, probably originated by the coalescence of “mini poles”, Asp-GFP also localizes to the divergent MT bundles and accumulates at their ends. Panels A5 and A6 illustrate how the divergent ends eventually coalesce giving rise to a bipolar spindle with Asp-GFP enriched at the poles. Scale bar, 5 µm. **B**. S2 cells expressing both Asp-GFP and Cherry-tubulin were fixed and stained with anti-α-tubulin (red), anti-GFP (green) and anti-Cid (white) antibodies and counterstained for DNA (DAPI, blue). Representative examples of early (B1 and B2) and late (B3 and B4) stages of spindle regrowth after colcemid-induced MT depolymerization illustrating the relationships between Asp-GFP, tubulin and centromeres (marked by Cid). Note that in all cases the Asp-GFP signals are surrounded by the centromeres, indicating that the MT clusters/asters are in fact generated by the coalescence of the minus ends of MT bundles emanating from the kinetochores. Scale bar, 5 µm.

Like Asp, Mast-GFP was accumulated on kinetochores at time 0, when KDMTR was not detectable (Figure 5, panel 1), Thus, Mast-GFP remains associated with kinetochores throughout the process of MT depolymerization. This suggests that Mast (CLASP1) is anchored to the kinetochore by some kinetochore-bound proteins. Although there is no experimental evidence supporting this occurrence, studies in human cells and *C. elegans* have shown that the CLASP proteins physically interact with CENPE and CENPF orthologs, respectively [74, 75]). Mast-GFP accumulation at the kinetochores persisted throughout the process of spindle reassembly, and Mast-GFP signals surrounded the MT clusters/asters (Figure 5). Mast-GFP also associated with the regrowing MT bundles, which, however, showed a much lower GFP fluorescence intensity than the kinetochores (Figure 5).

**Figure 5.**
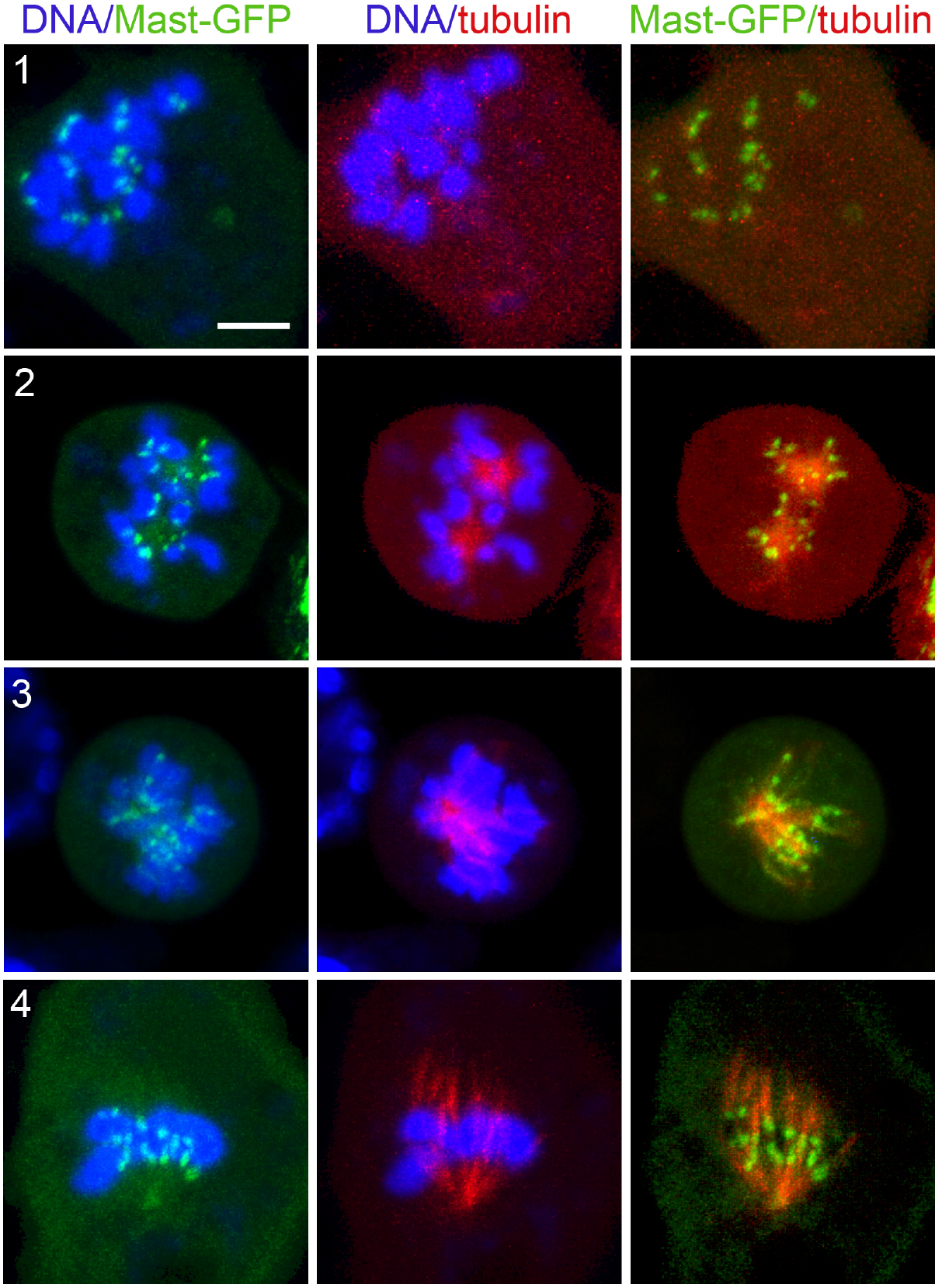
Mast-GFP localization during KDMTR after colcemid-induced MT depolymerization. Live S2 cells expressing Mast-GFP (green) and Cherry-tubulin (red) were stained with the vital DNA stain Hoechst 33342 (blue). Mast-GFP is associated with discrete sites on the chromosomes (probably corresponding to the kinetochores) after 3 h colcemid treatment, before washout of the drug and MTR (panel 1). During MTR, Mast-GFP is still accumulated on the kinetochores, which surround tubulin clusters (likely mini spindle poles) that exhibit weak GFP fluorescence (panel 2). Mast-GFP remains concentrated on kinetochores during further MTR stages (panels 3 and 4). Scale bar, 5 µm.

When spindle was completely depolymerized (time 0), Mars-GFP, Mei-38-GFP and Eb1-GFP were not associated with kinetochores. Mars-GFP and Mei-38-GFP co-localized with the initial MT foci at the beginning of spindle regrowth and remained on the spindle until it was completely reformed (Figure 6). Eb1-GFP was also permanently co-localized with the regrowing spindle structures. We filmed Eb1-GFP behavior at early regrowth stages, but we did not see the dynamic MT plus end-accumulations (comets) that are typically observed in normal PRO-METs. Regrowing MT structures were instead showing very small Eb1-GFP puncta that appeared to move along short paths both away and towards the kinetochores. However, completely, or almost completely reformed spindles displayed comets moving from the poles towards the spindle equator, like in untreated Eb1-GFP expressing cells (S1-S3 Movies). These observations are consistent with previous work on acentrosomal spindles assembled in Cnn-depleted cells [15], and suggest that, once formed, the poles of the acentrosomal spindles can drive MT formation.

**Figure 6.**
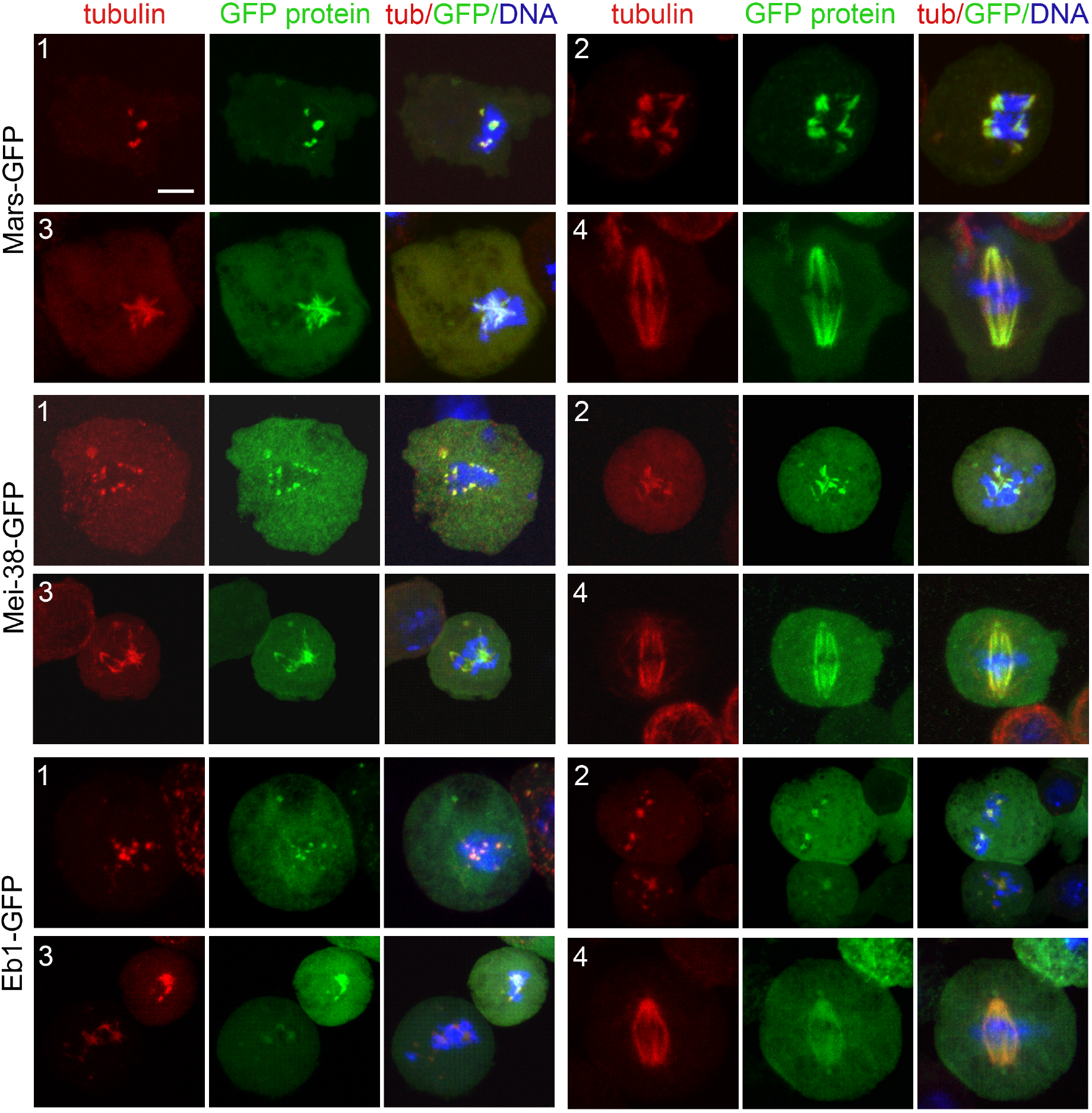
Localization of Mars-GFP, Mei-38-GFP and Eb1-GFP during KDMTR after colcemid-induced MT depolymerization. Fixed S2 cells expressing the indicated GFP-tagged protein were stained with anti-GFP antibodies (green), anti-α-tubulin antibodies (red) and DNA (DAPI, blue). Panels 1 show the initial MTR phases with small chromosome-associated tubulin signals that colocalize with the GFP proteins. Panels 2 and 3 show more advanced MTR phases characterized by the presence of MT asters/clusters. Panels 4 show reformed spindles. Note that in all cases the tubulin signals colocalize with the GFP signals. Scale bar, 5 µm.

As pointed out earlier, in untreated cells, the fluorescence of spindle-associated Patronin-GFP can either be dull or bright. Specifically, Patronin-GFP is bright when it moves from the kinetochores to the spindle poles along the k-fibers [53]. Here we considered only the bright fluorescence, as the dull one is difficult to score. PRO-METs at time 0 or at the very early MTR stages did not show any bright Patronin-GFP signal. These signals were seen in many MT clusters/asters, but their shapes were very different from those produced by Asp-GFP. They appeared as short bars within the MT clusters and were often associated with the MT bundles emanating from the clusters (Figure 7). These observations suggest that the bright form of Patronin-GFP marks the kinetochore-driven MTs when they start forming bundles and moves along these bundles as they keep growing.

**Figure 7.**
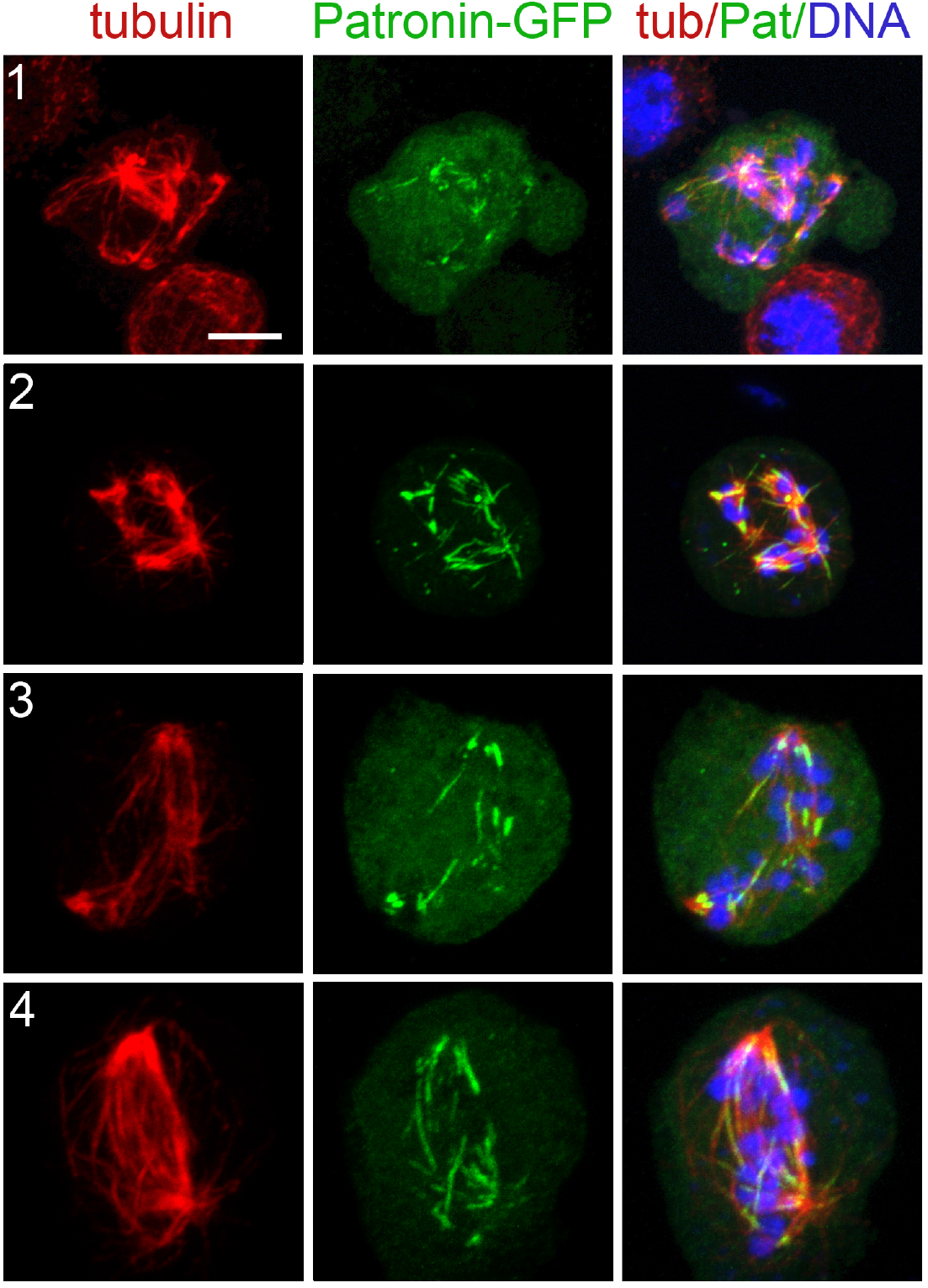
Patronin-GFP localization during KDMTR after colcemid-induced MTs depolymerization. S2 cells expressing both Patronin-GFP and Cherry-tubulin were fixed and stained with anti-α-tubulin (red) and anti-GFP (green) antibodies and counterstained for DNA (DAPI, blue). In very early MTR stages the highly fluorescent form of Patronin-GFP is not visible (see text). At later MTR stages, characterized by MT clusters/asters formation, Patronin-GFP is observed within these structures (panels 1 and 2) where it localizes along some but not all MT bundles. Localization along a fraction of the MT bundles is also seen in advanced stages of spindle reformation (panels 3 and 4). Scale bar, 5 µm.

## Discussion

### Genetic control of MT regrowth after tubulin depolymerization

We have shown that the mechanisms underlying KDMTR following colcemid-induced MT depolymerization can be genetically dissected using RNAi. In *mast, mei-38, mars, Eb1* and *dgt6* RNAi cells KDMTR is strongly reduced compared to control. In Patronin-depleted cells the advanced regrowth figures (elongated MT bundles and MT clusters) are slightly but not significantly less frequent than in control cells, but the mean CATF is significantly lower than in control. This finding could reflect a minor role of Patronin in KDMTR but could also be a consequence the limited efficiency of RNAi against *Patronin*. In our experimental conditions, *Patronin* RNAi cells showed 28% of residual mRNA, while the residual mRNAs for the other genes ranged from 2.8 to 8.4% of the control level (Table S1). In contrast, depletion of either Klp10A or Asp promotes KDMTR.

Previous work has suggested that in S2 cells lacking functional centrosomes kinetochore-driven MT formation is not controlled by the RCC1-RanGTP pathway [27]. Nonetheless, we found that the *Drosophila* homologs of TPX2 (Mei-38) and HURP (Mars) are required for KDMTR just as their vertebrate counterparts that are regulated by RanGTP [21, 32]. The *Xorbit* gene, the *Xenopus* homolog of *mast* (*CLASP1*), has also been shown to be required for chromatin-induced MTs nucleation [76], but there is no published evidence that the CLASP proteins control kinetochore-dependent MT formation in mammalian cells. To the best of our knowledge the possible roles of the vertebrate homologs of Eb1, Asp and Klp10A in KDMTR have never been addressed. However, given that at least Eb1, Asp and Klp10A appear to play conserved mitotic functions, it is quite possible that their vertebrate orthologues also control KDMTR.

We unexpectedly found that after colcemid-induced MT depolymerization, control and RNAi cells display very similar KDMTR figures and that the main difference between control and RNAi samples was in the frequency of these figures at different fixation times (Figure 2, Table 2). Thus, it appears that our RNAi treatments either delay or stimulate KDMTR without affecting the pattern of MTR. This observation points to a particular genetic robustness of the KDMTR process and suggests that it is under the control of many genes that are at least in part functionally redundant. This is not surprising, given that an efficient cell division is at the basis of life and that the kinetochore-driven MT formation pathway is not only necessary but also sufficient for spindle assembly in most eukaryotic cells.

### Kinetochore-driven MT growth and spindle length

Our results indicate that the efficiency of KDMTR in RNAi cells positively correlates with the length of the spindles observed in the same cells. For example, the spindles of Mast-, Mei-38-, Mars-, Dgt6-, Patronin- and Eb1-depleted cells are significantly shorter than control spindles, while the spindles of *asp* RNAi cells are longer than in control (Figure S1; see also [23, 50-53]. Also the spindles of Klp10A-depleted cells are longer than control spindles; the increased KDMTR and spindle length in these cells are a likely consequence of a reduced Klp10A-mediated MT depolymerization [50, 52]. Other *Drosophila* proteins that promote KDMTR and whose loss results in short spindles are Misato (Mst) and its interacting partners of the TCP-1 and Prefoldin complexes [33, 34], Ensconsin that shares homology with human MAP7 [35] and the Msps (TOGp) MT polymerase [23, 50]. Thus, at least in *Drosophila* S2 cells KDMTR effectiveness strongly correlates with the length of prometaphase/metaphase spindles. RNAi-based screens in S2 cells identified many additional genes associated with a short spindle phenotype [16, 65] but whether their loss results in decreased KDMTR is currently unknown.

### Kinetochore-driven MT growth during normal mitosis and after MT depolymerization

One of the main questions is whether and to which extent KDMTR recapitulates the MT growth that occur at kinetochores during normal cell division. As mentioned earlier, centrosome-containing S2 cells were used for a pivotal analysis of kinetochore-driven MT growth. Maiato and coworkers (2004) [20] showed that living cells stably expressing GFP-α-tubulin form k-fibers from the unattached kinetochore of mono-oriented chromosomes and that these fibers extend towards the cell periphery until they interact with the astral MTs and become oriented towards the spindle poles. They also used laser beam microsurgery to show that when individual k-fibers are severed, the fragment associated with the kinetochore (k-fragment) regrows, while the fragment terminating in the spindle pole degenerates. Finally, using an elegant combination of laser microsurgery/photobleaching, they were able to demonstrate that the growth of the k-fragment starts at or near the kinetochore. Based on these data, they proposed that kinetochore-driven MT formation begins with the generation of short randomly oriented MTs near the kinetochores. The plus ends of these MTs would then be captured by the kinetochore and continue to polymerize there, leading to k-fiber elongation. As a result, the MT minus ends would be pushed away from the kinetochores and accumulate at the end of the regrowing k-fiber [20].

One aspect of this model that has been matter of debate is the mechanism underlying the initial MTR. Mishra and coworkers [38] found that Nup107-160 and γ-tubulin localize next to the kinetochores and tubulin regrowth foci in HeLa cells at the early MTR stages after nocodazole-induced MT depolymerization. They also showed that KDMTR is reduced in the absence of either Nup107-160 or γ-tubulin. They concluded that although these results are compatible with the Maiato’s model [20], they are also consistent with the possibility that during the initial stages of KDMTR MTs are anchored to the kinetochores through their minus ends and polymerize at the distal plus ends, forming an assembly intermediate that disappear as k-fibers elongate [38]. A similar transient polarity inversion of kinetochore MTs has been described in budding yeast, where kinetochores are thought to generate MTs with the minus ends embedded into the kinetochores and the plus ends pointing outwards; these MTs disappear when the plus ends of the MTs that extend from the spindle poles are captured by the kinetochores and do not contribute to the metaphase spindle [77].

We have shown that Asp-GFP accumulates at kinetochores when MTR is not detectable and then partially co-localizes with the first-kinetochore-associated small foci of regrowing MTs. As MTR proceeds, Asp-GFP is consistently found at the center of MT asters/clusters regardless of their size. These structures do not exhibit kinetochores at their centers but are instead surrounded by kinetochores. This suggests that Asp-GFP binds the minus ends of the regrowing MT bundles since the beginning of their formation and that the minus ends of these bundles eventually converge giving rise to a sort of mini spindle poles with Asp-GFP at their center. Asp-GFP then localizes on the long MT bundles/k-fibers that emanate from these asters/clusters and accumulates at their extremities, indicating that they are indeed enriched in MT minus ends [20]. The simplest interpretation of these observations is that Asp-GFP associates with the MT minus ends emanating from kinetochores and is pushed away by the growth of the MT plus ends embedded into these organelles. Thus, our data argue against the possibility of a transient MT polarity inversion during the early stages of MTR after colcemid-induced depolymerization.

Recent ultrastructural studies revealed that most kinetochores of unperturbed human cells in very early prometaphase are transiently associated with a mesh of short randomly oriented noncentrosomal MTs, which is no longer observed upon formation of mature k-fibers [25]. Although there is no direct evidence that this MT mesh is present also in early S2 cell prometaphases, our data are consistent with its existence at least in cells undergoing KDMTR. It is likely that Asp-GFP binds the minus ends of the short MTs that form near the centromeres, limiting their growth, as suggested by the finding that in Asp-depleted cells KDMTR is increased compared to control. In human cells, ASPM forms a complex with katanin, and the two proteins cooperate in promoting MT severing [58]. Whether katanin binds *Drosophila* Asp is unknown, but if so, katanin might contribute to the regulation of MT minus ends also during S2 cells mitosis. The finding that during KDMTR Asp is transiently associated with the kinetochores further suggests that this protein might have role in the establishment of a correct kinetochore-MT attachment and explains why Asp-depleted cells suffer a spindle checkpoint-dependent metaphase arrest [78].

### An integrated model on kinetochore-driven MT growth

Our results provide insight into the genetic/molecular control of KDMTR. Previous work on *Drosophila* S2 cells has shown that Mast mediates the incorporation of tubulin subunits into the plus ends of the mature k-fibers embedded into the kinetochore [61]. It has been also suggested that human CLASP1 plays a similar function [79]. Our findings that depletion of Mast results in a strong reduction of KDMTR and that Mast-GFP co-localizes with the kinetochores in all phases of MTR strongly suggest that this protein is the main determinant of MT plus end elongation during KDMTR.

We found that Mars (HURP), Mei-38 (TPX2) and the augmin subunit Dgt6 are required for KDMTR. Studies in vertebrate systems have shown that TPX2 interacts with a variety of mitotic proteins, including Aurora A (AURKA) and the augmin complex, promotes MT nucleation and bundling, and is required for both kinetochore fiber formation and KDMTR [3, 21, 80]. HURP has MT bundling activity and is required for k-fiber stability and KDMTR [22, 81]. Augmin is thought to promote MT nucleation within the k-fibers by recruiting the γ-TURCs (reviewed in [3]). We have shown that these proteins precisely co-localize with the kinetochore-driven MTs from the very early stages of regrowth to the complete reformation of the spindles. This suggests that these three proteins play conserved functions required for proper reassembly and stability of k-fibers after MT depolymerization.

We also found that KDMTR is negatively affected by loss of Eb1 and at least in part by Patronin depletion. The two proteins showed very different patterns of localization during KDMTR. Eb1-GFP localized to the kinetochore-dependent MTs from the beginning of repolymerization and remained on these MTs until the spindle was reassembled. In contrast, we observed the brightly fluorescent form of Patronin-GFP only when the kinetochores started to emanate MT bundles. Our analyses on live and fixed cells also showed that brightly fluorescent Patronin-GFP is highly dynamic and tends to move from the kinetochores to the spindle poles in unperturbed S2 cells [53]. We speculate that Patronin-GFP moves along the regrowing MT bundles exploiting an as yet unidentified minus-end-directed motor. During its movement, Patronin-GFP would associate with the MT minus ends including those generated by the augmin pathway. This would help maintaining the correct structural organization and stability of mitotic MT bundles [53]. Notably, the human homologs of Patronin (CAMSAP2, CAMSAP1, CAMSAP3) bind katanin just like ASPM [82] but it is currently unknown whether this occurs also in *Drosophila*. In any case, our results suggest that loss of Patronin does not result in reduced severing of spindles MTs.

In summary, our results indicate that KDMTR after MT depolymerization recapitulates the process of kinetochore-driven MT formation in unperturbed cells. Our results also suggest an integration of the current model for kinetochore-dependent MT formation in *Drosophila* S2 cells [20]. We propose that kinetochores capture the plus ends of MTs nucleated in their vicinity and that these MTs elongate through the action of Mast. These processes are likely to be downregulated by Asp that binds the MT minus ends and prevents excessive and disorganized MTR. Mars, Mei-38, Dgt6, Eb1 and Patronin positively regulate the subsequent formation, elongation and stabilization of the regrowing MT bundles/k-fibers.

## Materials and Methods

### Cell cultures

The S2 cells used for colcemid-induced MT depolymerization were grown 39.4 g/L Shields and Sang M3 Insect medium (Sigma) supplemented with 0.5 g/L KHCO_3_ and 20% heat-inactivated FBS (Thermo Fisher Scientific). S2 cells expressing GFP-tagged proteins, including cells expressing GFP-tubulin (obtained from DGRC) [64], were cultured in 39.4 g/L Shields and Sang M3 Insect medium (Sigma) supplemented with 2.5 g/L bacto peptone (Difco), 1 g/L yeast extract (Difco), and 5% heat-inactivated FBS (Thermo Fisher Scientific). All cells were cultured at 25°C and were free from mycoplasma contamination.

### dsRNA production and RNAi treatments

The DNA templates for the synthesis of dsRNAs for RNAi against *mast, mei-38, mars, dgt6, Eb1, Patronin, asp* and *Klp10A* were amplified from wild type (*Oregon-R*) genomic DNA or cDNA libraries made from 0-24 h *Oregon-R* embryos. The primer sequences used for amplification are reported in Table S2. The PCR products were purified using spin columns. Synthesis of dsRNAs was carried out as described earlier [83].

RNAi treatments were carried out as follows. 1×10^6^ S2 cells in 1 ml of the appropriate serum-free medium were plated in a well of a six-well culture dish. 60-80 µg of dsRNA was then added to each well. After 1 h incubation, 2 ml of the FBS-supplemented medium was added to the wells, and cells were grown for 5 days. For *Patronin* and *Klp10A* RNAi, a second dose (60-80 µg) of dsRNA was added to the samples on day 3, and cells were grown for 2 additional days. Control S2 cells were prepared in the same way, but without addition of dsRNA.

### Evaluation of the RNAi efficiency by RT-qPCR

Gene-specific primers (different from those used for ds RNA production) were designed using Primer-BLAST (https://www.ncbi.nlm.nih.gov/tools/primer-blast/) or Primer3 (http://bioinfo.ut.ee/primer3-0.4.0/primer3/) software; primer sequences are provided in Table S3. For each primer pair, the amplification efficiency was determined from the slope of the log-linear portion of the calibration curve [84] generated using dilutions of cDNA prepared from wild-type S2 cells (Table S3). Total RNA was isolated from control and dsRNA-treated S2 cells using RNAzol® RT reagent (MRC) according to the manufacturer’s instructions. Reverse transcription was performed with the RevertAid reverse transcriptase (Thermo Fisher Scientific) using 2 µg of total RNA in the presence of 2 U/µl of RNaseOut Recombinant RNase Inhibitor (Thermo Fisher Scientific). Genomic DNA was eliminated using the RapidOut DNA Removal Kit (Thermo Fisher Scientific). qPCR was carried out using BioMaster HS-qPCR SYBR Blue (2×) reagent kit (Biolabmix; https://biolabmix.ru/en/) and CFX96™ Real-Time PCR Detection System (Bio-Rad). We used the following thermal cycling conditions: 5 min at 95°C, followed by 39 cycles of 15 sec at 95°C, 30 sec at 60°C, and 30 sec at 72°C. Data were collected during each extension phase. Negative control templates (water and cDNA synthesized without reverse transcriptase) were included in each run. Measurements of gene expression were done at least in two biological replicates, each with three technical replicates. The relative mRNA quantification was determined using the ΔΔCq method [84]. mRNA expression levels were normalized to those of the housekeeping gene *RpL32*; primers for this gene are from [85]).

### Colcemid-induced MT depolymerization

Control and RNAi samples were incubated in a medium containing colcemid (Calbiochem, Merck Millipore) at the final concentration of 4.5 µg/ml. After 3 h incubation, cells were centrifuged (1000 *g*, 22°C, 5 min) and the colcemid-containing medium was removed. Cells were washed three times in drug-free medium and left in this medium. Cells were then fixed after 20-75 min after colcemid removal or handed over for *in vivo* observation by confocal microscopy (cells expressing GFP-tagged proteins). A part of the cells was fixed at the end of 3 h colcemid treatment (without removal of the drug) to check the degree of MT depolymerization (time 0).

### Generation of S2 cells expressing GFP-tagged proteins

S2 cell lines expressing mCherry-αTub84B (referred to as Cherry-tubulin throughout the manuscript) and either Asp-GFP or Patronin-GFP, were generated previously [53]. The other cell lines expressing both Cherry-tubulin and GFP-tagged proteins used here were generated using a piggyBac transposon-based vector as described in [53, 83]. Full-length coding sequences of *Eb1, mars, mast* and *mei-38* were cloned in the piggyBac transposon vector between the copper-inducible *MtnA* promoter and the enhanced GFP (referred to as GFP throughout the manuscript) coding sequence. The resulting plasmid constructs, verified by sequencing, encode the C-terminal GFP fusions of Eb1 (GenPept accession no. AGB93267.1), Mars (GenPept accession no. AHN56186.1), Mast (GenPept accession no. AAN12151.1), or Mei-38 (GenPept accession no. AAF45636.1, but with V99D, A117D, S120P and E162_R163insA mutations, caused by known DNA sequence variations (http://dgrp.gnets.ncsu.edu/; [86, 87]). Each plasmid also contained the Cherry-tubulin coding sequence under the control of the *MtnA* promoter, and a blasticidin-resistance cassette. Details of the plasmid constructions are available upon request. Integration of the transgenes in the genome of S2 cells and the subsequent selection of modified cells with blasticidin were done as described in [83]. All cells were free from mycoplasma contamination. To induce expression of fluorescent fusion proteins, cells were grown in the presence of 100-250 µM of copper sulfate for 16-22 h before *in vivo* analysis or fixation.

### Fixation, immunostaining and microscope analysis

For mitotic phenotype analysis cells were fixed as described in [88]. Briefly, 2×10^6^ S2 cells were spun down by centrifugation at 1000 *g* for 5 min, washed in 2 ml of 1×PBS (Sigma), and fixed for 10 min in 2 ml of 3.7% formaldehyde in 1×PBS. Cells were resuspended in 500 µl of PBS, and cytocentrifuged on clean slides (using a Cytospin™ 4 Cytocentrifuge, Thermo Fisher Scientific, at 900 rpm for 4 min). Slides were then immersed in liquid nitrogen for 5 min, transferred to PBT (PBS plus 0.1% TritonX-100) for 30 min, and then to PBT containing 3% BSA (AppliChem) for 30 min.

The slides were immunostained using the following primary antibodies, all diluted in a 1:1 mixture of PBT and 3% BSA: mouse anti-α-tubulin (1:500, Sigma T6199), rabbit anti-Spd-2 (1:4.000, [14]), rabbit anti-Cid (1:300, Abcam ab10887) and chicken anti-GFP (1:200, Invitrogen PA1-9533). Primary antibodies were detected by incubation for 1 h with FITC-conjugated goat anti-mouse IgG (1:40, Sigma F8264) or Alexa Fluor 568-conjugated goat anti-mouse IgG (1:300, Invitrogen A11031), Alexa Fluor 568-conjugated goat anti-rabbit IgG (1:350, Invitrogen A11036) or Alexa Fluor 660-conjugated goat anti-rabbit IgG (1:300, Invitrogen A21074), and Alexa Fluor 488-conjugated goat anti-chicken IgG (1:300, Invitrogen A11039). To reduce fluorescence fading most slides were mounted in Vectashield antifade mounting medium containing 4.6-diamidino-2-phenylindole (DAPI; Vector Laboratories). Slides stained with Alexa Fluor 660 were stained with DAPI and then mounted in the ProLongTM Gold Antifade Mountant (Thermo Fisher Scientific). Images of fixed cells were captured using a Zeiss Axio Imager M2 equipped with an EC Plan-Neofluar 100×/1.30 oil lens (Carl Zeiss) and with an AxioCam 506 mono (D) camera. Some of the fixed cell images were generate using a Zeiss LSM 710 confocal microscope and an oil immersion 100×/1.40 plan-apo lens with an AxioCam 506 mono camera using the ZEN 2012 software (Carl Zeiss).

For live cell analysis, 500 µl of cell suspensions (5×10^5^ cells/ml) were transferred to cell chambers (Invitrogen A-7816) containing coverslips treated with 0.25mg/ml concanavalin A (Sigma-Aldrich C0412) placed on the bottom of the chambers. In some cases, live cells were treated with Hoechst 33342 (Invitrogen H3570, final concentration 20 μg/ml) to stain chromosomal DNA. Observations were performed between 20 and 180 min after cell plating using the same Zeiss confocal microscope described above.

### Spindle length measurements

The spindle length was measured with the ZEN 2012 software (Carl Zeiss) as described in detail in [83]. Measures were restricted to cells that did not appear to be polyploid with respect to the basic karyotype of S2 cells. The data were compared using the Mann–Whitney *U* test and plotted using the BoxPlotR web-tool (http://shiny.chemgrid.org/boxplotr/) [89].

### Fluorescence intensity measures

The fluorescence intensity of chromosome-associated regrowing MTs after colcemid-induced MT depolymerization was measured with the ImageJ program [90] using the “Freehand” selection tool and the function “Measure”. In all cases, we selected the tubulin signals that were clearly associated with the kinetochores/chromosomes and measured them individually. To calculate the CATF in a cell, we subtracted the background fluorescence from the fluorescence of each tubulin signal. To make comparable different experiments, we then normalized the corrected fluorescence values of both RNAi and control cells from the same biological replicate to the mean fluorescence value of the control.

## Acknowledgments

We thank Maria B. Schwartz (Berkaeva) for her help during the early stages if this work. We also thank Mikhail O. Lebedev and Tatiana D. Dubatolova for their technical assistance. DNA sequencing and confocal microscopy analysis were performed using resources provided by the Molecular and Cellular Biology core facility of the Institute of Molecular and Cellular Biology SB RAS.

## Supporting information

**S1 Table. RNAi efficiency and frequencies of mitotic figures observed after RNAi against *asp, dgt6, Eb1, Klp10A, mars, mast, mei-38* and *Patronin***.

**S2 Table. dsRNAs used for RNA interference**.

**S3 Table. Primers used for assessment of the RNAi-mediated gene silencing by RT-qPCR**

**S1 Figure. Spindle length of PRO-METs in untreated and RNAi S2 cells**. Consistent with previous results, the spindles of *mast, mars, mei-38, Eb1* and *Patronin* RNAi cells are significantly shorter than control spindles, while those of *asp* and *Klp10A* RNAi cells are significantly longer. The red line represents the median of the control length. * P < 0.05; Mann–Whitney *U* test.

**S2 Figure. A representative example of spindle reformation after colcemid-induced MT depolymerization**. The live S2 cell shown, which expresses GFP-tubulin, was stained with the vital DNA stain (Hoechst 33342 (blue); shown only at time 0:00). Timescale, h:min. Note that at the beginning of imaging (time 0:00) there are only a few small tubulin regrowth foci associated with the chromosomes. These foci grow forming tubulin clusters (time 0:07) that coalesce producing a spindle that is focused at only one of the poles (time 0:22). The MT bundles at the unfocused pole progressively converge forming a second pole (time 0:32-0:59). Scale bar, 5 µm.

**S3 Figure. Frequencies per cell of MT regrowth figures in RNAi and control cells observed 30 min after colcemid removal**. Data are from Table 2. Short, MT spots and short bundles; long, elongated MT bundles; cluster. MT clusters/asters (see Figures 1 and 2). Data have been analyzed with the Mann-Whitney *U* test; ns, not significant; **, ***, **** significant with p < 0.01, 0.001, and 0.0001, respectively. Error bars SEM.

**S4 Figure. Relative CATF observed in the indicated RNAi cells and control cells**. Row fluorescence measures of each biological replicate (RNAi cells and pertinent control cells) were normalized to the mean fluorescence of control cells and compared using the Mann-Whitney *U* test; ns, not significant; *, **, ***, **** significant with p < 0.05, 0.01, 0.001, and 0.0001, respectively. Error bars SEM.

**S5 Figure. Mast-GFP localization during mitosis of S2 cells**. Live interphase and mitotic S2 cells expressing both Mast-GFP and Cherry-tubulin. Note that Mast-GFP accumulates at the kinetochores, the centrosomes and the central spindle of telophase cells. Scale bar, 5 µm.

**S6 Figure. Mei-38-GFP localization during mitosis of S2 cells**. Live interphase and mitotic S2 cells expressing both Mei-38-GFP and Cherry-tubulin. Note that Mei-38-GFP associates with the spindle MTs throughout mitosis. Scale bar, 5 µm.

**S7 Figure. Mars-GFP localization during mitosis of S2 cells**. Live interphase and mitotic S2 cells expressing both Mars-GFP and Cherry-tubulin. Note that Mars-GFP accumulates in the nucleolus in interphase and late telophase cells. During metaphase and early anaphase, Mars-GFP associates with the spindle MTs but is excluded from the centrosome area. In late anaphase and telophase, Mars-GFP is no longer associated with the spindle MTs. Scale bar, 5 µm.

**S8 Figure. Eb1-GFP localization during mitosis of S2 cells**. Live interphase and mitotic S2 cells expressing both Eb1-GFP and Cherry-tubulin. Note that Eb1-GFP associates with the spindle throughout mitosis showing accumulation at the MT plus ends. Scale bar, 5 µm.

**S9 Figure. Asp-GFP localization during mitosis of S2 cells**. Live interphase and mitotic S2 cells expressing both Asp-GFP and Cherry-tubulin. Note the Asp-GFP accumulation in the interphase nucleus, at the spindle poles and at the MT minus end-enriched extremities of the telophase central spindle. Scale bar, 5 µm.

**S10 Figure. A representative example of spindle reformation after colcemid-induced MT depolymerization in S2 cells expressing both Asp-GFP and Cherry-tubulin**. The live S2 cell shown, which expresses both Asp-GFP (green) and Cherry-tubulin (red), was stained with the vital DNA stain (Hoechst 33342 (blue); shown only at time 0:00). Timescale, h:min. Note that at the beginning of imaging (time 0:00) Asp-GFP localizes at the center of MT clusters/asters. Asp-GFP remains in its initial location that defines one of the spindle poles but also moves to the ends of the regrowing MT bundles to form a second spindle pole.

**S1 Movie. Microtubule dynamics in an unperturbed S2 cell prometaphase spindle**. The cell expressing Eb1-GFP is stained with DNA vital dye Hoechst 33342.

**S2 Movie. Microtubule dynamics in a MT cluster during KDMTR after colcemid-induced MT depolymerization**. The S2 cell expressing Eb1-GFP is stained with DNA vital dye Hoechst 33342.

**S3 Movie. Microtubule dynamics in an almost completely reformed prometaphase spindle following colcemid-induced MT depolymerization**. The S2 cell expressing Eb1-GFP is stained with DNA vital dye Hoechst 33342.

## References

1. O’Connell CB, Khodjakov AL. Cooperative mechanisms of mitotic spindle formation. J Cell Sci. 2007;120(Pt 10):1717–22. https://doi.org/10.1242/jcs.03442. PMID: 17502482.

2. Duncan T, Wakefield JG. 50 ways to build a spindle: the complexity of microtubule generation during mitosis. Chromosome Res. 2011;19(3):321–33. https://doi.org/10.1007/s10577-011-9205-8. PMID: 21484448.

3. Petry S. Mechanisms of Mitotic Spindle Assembly. Annu Rev Biochem. 2016;85:659–83. https://doi.org/10.1146/annurev-biochem-060815-014528. PMID: 27145846.

4. Prosser SL, Pelletier L. Mitotic spindle assembly in animal cells: a fine balancing act. Nat Rev Mol Cell Biol. 2017;18(3):187–201. https://doi.org/10.1038/nrm.2016.162. PMID: 28174430.

5. Luders J, Stearns T. Microtubule-organizing centres: a re-evaluation. Nat Rev Mol Cell Biol. 2007;8(2):161–7. https://doi.org/10.1038/nrm2100. PMID: 17245416.

6. Gatti M, Bucciarelli E, Lattao R, Pellacani C, Mottier-Pavie V, Giansanti MG, et al. The relative roles of centrosomal and kinetochore-driven microtubules in Drosophila spindle formation. Exp Cell Res. 2012;318(12):1375–80. https://doi.org/10.1016/j.yexcr.2012.05.001. PMID: 22580224.

7. Meunier S, Vernos I. Acentrosomal Microtubule Assembly in Mitosis: The Where, When, and How. Trends Cell Biol. 2016;26(2):80–7. https://doi.org/10.1016/j.tcb.2015.09.001. PMID: 26475655.

8. Walczak CE, Heald R. Mechanisms of mitotic spindle assembly and function. Int Rev Cytol. 2008;265:111–58. https://doi.org/10.1016/S0074-7696(07)65003-7. PMID: 18275887.

9. Khodjakov A, Cole RW, Oakley BR, Rieder CL. Centrosome-independent mitotic spindle formation in vertebrates. Curr Biol. 2000;10(2):59–67. https://doi.org/10.1016/s0960-9822(99)00276-6. PMID: 10662665.

10. Bonaccorsi S, Giansanti MG, Gatti M. Spindle assembly in Drosophila neuroblasts and ganglion mother cells. Nat Cell Biol. 2000;2(1):54–6. https://doi.org/10.1038/71378. PMID: 10620808.

11. Megraw TL, Kao LR, Kaufman TC. Zygotic development without functional mitotic centrosomes. Curr Biol. 2001;11(2):116–20. https://doi.org/10.1016/s0960-9822(01)00017-3. PMID: 11231128.

12. Basto R, Lau J, Vinogradova T, Gardiol A, Woods CG, Khodjakov A, et al. Flies without centrioles. Cell. 2006;125(7):1375–86. https://doi.org/10.1016/j.cell.2006.05.025. PMID: 16814722.

13. Blachon S, Gopalakrishnan J, Omori Y, Polyanovsky A, Church A, Nicastro D, et al. Drosophila asterless and vertebrate Cep152 Are orthologs essential for centriole duplication. Genetics. 2008;180(4):2081–94. https://doi.org/10.1534/genetics.108.095141. PMID: 18854586.

14. Giansanti MG, Bucciarelli E, Bonaccorsi S, Gatti M. Drosophila SPD-2 is an essential centriole component required for PCM recruitment and astral-microtubule nucleation. Curr Biol. 2008;18(4):303–9. https://doi.org/10.1016/j.cub.2008.01.058. PMID: 18291647.

15. Mahoney NM, Goshima G, Douglass AD, Vale RD. Making microtubules and mitotic spindles in cells without functional centrosomes. Curr Biol. 2006;16(6):564–9. https://doi.org/10.1016/j.cub.2006.01.053. PMID: 16546079.

16. Somma MP, Ceprani F, Bucciarelli E, Naim V, De Arcangelis V, Piergentili R, et al. Identification of Drosophila mitotic genes by combining co-expression analysis and RNA interference. PLoS Genet. 2008;4(7):e1000126. https://doi.org/10.1371/journal.pgen.1000126. PMID: 18797514.

17. Moutinho-Pereira S, Debec A, Maiato H. Microtubule cytoskeleton remodeling by acentriolar microtubule-organizing centers at the entry and exit from mitosis in Drosophila somatic cells. Mol Biol Cell. 2009;20(11):2796–808. https://doi.org/10.1091/mbc.E09-01-0011. PMID: 19369414.

18. Hayward D, Metz J, Pellacani C, Wakefield JG. Synergy between multiple microtubule-generating pathways confers robustness to centrosome-driven mitotic spindle formation. Dev Cell. 2014;28(1):81–93. https://doi.org/10.1016/j.devcel.2013.12.001. PMID: 24389063.

19. Witt PL, Ris H, Borisy GG. Origin of kinetochore microtubules in Chinese hamster ovary cells. Chromosoma. 1980;81(3):483–505. https://doi.org/10.1007/BF00368158. PMID: 7449572.

20. Maiato H, Rieder CL, Khodjakov A. Kinetochore-driven formation of kinetochore fibers contributes to spindle assembly during animal mitosis. J Cell Biol. 2004;167(5):831–40. https://doi.org/10.1083/jcb.200407090. PMID: 15569709.

21. Tulu US, Fagerstrom C, Ferenz NP, Wadsworth P. Molecular requirements for kinetochore-associated microtubule formation in mammalian cells. Curr Biol. 2006;16(5):536–41. https://doi.org/10.1016/j.cub.2006.01.060. PMID: 16527751.

22. Torosantucci L, De Luca M, Guarguaglini G, Lavia P, Degrassi F. Localized RanGTP accumulation promotes microtubule nucleation at kinetochores in somatic mammalian cells. Mol Biol Cell. 2008;19(5):1873–82. https://doi.org/10.1091/mbc.E07-10-1050. PMID: 18287525.

23. Bucciarelli E, Pellacani C, Naim V, Palena A, Gatti M, Somma MP. Drosophila Dgt6 interacts with Ndc80, Msps/XMAP215, and gamma-tubulin to promote kinetochore-driven MT formation. Curr Biol. 2009;19(21):1839–45. https://doi.org/10.1016/j.cub.2009.09.043. PMID: 19836241.

24. O’Connell CB, Loncarek J, Kalab P, Khodjakov A. Relative contributions of chromatin and kinetochores to mitotic spindle assembly. J Cell Biol. 2009;187(1):43–51. https://doi.org/10.1083/jcb.200903076. PMID: 19805628.

25. Sikirzhytski V, Renda F, Tikhonenko I, Magidson V, McEwen BF, Khodjakov A. Microtubules assemble near most kinetochores during early prometaphase in human cells. J Cell Biol. 2018;217(8):2647–59. https://doi.org/10.1083/jcb.201710094. PMID: 29907657.

26. Kalab P, Heald R. The RanGTP gradient - a GPS for the mitotic spindle. J Cell Sci. 2008;121(Pt 10):1577–86. https://doi.org/10.1242/jcs.005959. PMID: 18469014.

27. Moutinho-Pereira S, Stuurman N, Afonso O, Hornsveld M, Aguiar P, Goshima G, et al. Genes involved in centrosome-independent mitotic spindle assembly in Drosophila S2 cells. Proc Natl Acad Sci U S A. 2013;110(49):19808–13. https://doi.org/10.1073/pnas.1320013110. PMID: 24255106.

28. Chen JW, Barker AR, Wakefield JG. The Ran Pathway in Drosophila melanogaster Mitosis. Front Cell Dev Biol. 2015;3:74. https://doi.org/10.3389/fcell.2015.00074. PMID: 26636083.

29. Goshima G, Nedelec F, Vale RD. Mechanisms for focusing mitotic spindle poles by minus end-directed motor proteins. J Cell Biol. 2005;171(2):229–40. https://doi.org/10.1083/jcb.200505107. PMID: 16247025.

30. Kallio MJ, McCleland ML, Stukenberg PT, Gorbsky GJ. Inhibition of aurora B kinase blocks chromosome segregation, overrides the spindle checkpoint, and perturbs microtubule dynamics in mitosis. Curr Biol. 2002;12(11):900–5. https://doi.org/10.1016/s0960-9822(02)00887-4. PMID: 12062053.

31. Koffa MD, Casanova CM, Santarella R, Kocher T, Wilm M, Mattaj IW. HURP is part of a Ran-dependent complex involved in spindle formation. Curr Biol. 2006;16(8):743–54. https://doi.org/10.1016/j.cub.2006.03.056. PMID: 16631581.

32. Casanova CM, Rybina S, Yokoyama H, Karsenti E, Mattaj IW. Hepatoma up-regulated protein is required for chromatin-induced microtubule assembly independently of TPX2. Mol Biol Cell. 2008;19(11):4900–8. https://doi.org/10.1091/mbc.E08-06-0624. PMID: 18799614.

33. Mottier-Pavie V, Cenci G, Verni F, Gatti M, Bonaccorsi S. Phenotypic analysis of misato function reveals roles of noncentrosomal microtubules in Drosophila spindle formation. J Cell Sci. 2011;124(Pt 5):706–17. https://doi.org/10.1242/jcs.072348. PMID: 21285248.

34. Palumbo V, Pellacani C, Heesom KJ, Rogala KB, Deane CM, Mottier-Pavie V, et al. Misato Controls Mitotic Microtubule Generation by Stabilizing the TCP-1 Tubulin Chaperone Complex [corrected]. Curr Biol. 2015;25(13):1777–83. https://doi.org/10.1016/j.cub.2015.05.033. PMID: 26096973.

35. Gallaud E, Caous R, Pascal A, Bazile F, Gagne JP, Huet S, et al. Ensconsin/Map7 promotes microtubule growth and centrosome separation in Drosophila neural stem cells. J Cell Biol. 2014;204(7):1111–21. https://doi.org/10.1083/jcb.201311094. PMID: 24687279.

36. Meunier S, Vernos I. K-fibre minus ends are stabilized by a RanGTP-dependent mechanism essential for functional spindle assembly. Nat Cell Biol. 2011;13(12):1406–14. https://doi.org/10.1038/ncb2372. PMID: 22081094.

37. Meunier S, Shvedunova M, Van Nguyen N, Avila L, Vernos I, Akhtar A. An epigenetic regulator emerges as microtubule minus-end binding and stabilizing factor in mitosis. Nat Commun. 2015;6:7889. https://doi.org/10.1038/ncomms8889. PMID: 26243146.

38. Mishra RK, Chakraborty P, Arnaoutov A, Fontoura BM, Dasso M. The Nup107-160 complex and gamma-TuRC regulate microtubule polymerization at kinetochores. Nat Cell Biol. 2010;12(2):164–9. https://doi.org/10.1038/ncb2016. PMID: 20081840.

39. Goshima G, Mayer M, Zhang N, Stuurman N, Vale RD. Augmin: a protein complex required for centrosome-independent microtubule generation within the spindle. J Cell Biol. 2008;181(3):421–9. https://doi.org/10.1083/jcb.200711053. PMID: 18443220.

40. Petry S, Groen AC, Ishihara K, Mitchison TJ, Vale RD. Branching microtubule nucleation in Xenopus egg extracts mediated by augmin and TPX2. Cell. 2013;152(4):768–77. https://doi.org/10.1016/j.cell.2012.12.044. PMID: 23415226.

41. Song JG, King MR, Zhang R, Kadzik RS, Thawani A, Petry S. Mechanism of how augmin directly targets the gamma-tubulin ring complex to microtubules. J Cell Biol. 2018;217(7):2417–28. https://doi.org/10.1083/jcb.201711090. PMID: 29875259.

42. Lawo S, Bashkurov M, Mullin M, Ferreria MG, Kittler R, Habermann B, et al. HAUS, the 8-subunit human Augmin complex, regulates centrosome and spindle integrity. Curr Biol. 2009;19(10):816–26. https://doi.org/10.1016/j.cub.2009.04.033. PMID: 19427217.

43. Zhu H, Coppinger JA, Jang CY, Yates JR, 3rd, Fang G. FAM29A promotes microtubule amplification via recruitment of the NEDD1-gamma-tubulin complex to the mitotic spindle. J Cell Biol. 2008;183(5):835–48. https://doi.org/10.1083/jcb.200807046. PMID: 19029337.

44. Uehara R, Nozawa RS, Tomioka A, Petry S, Vale RD, Obuse C, et al. The augmin complex plays a critical role in spindle microtubule generation for mitotic progression and cytokinesis in human cells. Proc Natl Acad Sci U S A. 2009;106(17):6998–7003. https://doi.org/10.1073/pnas.0901587106. PMID: 19369198.

45. David AF, Roudot P, Legant WR, Betzig E, Danuser G, Gerlich DW. Augmin accumulation on long-lived microtubules drives amplification and kinetochore-directed growth. J Cell Biol. 2019;218(7):2150–68. https://doi.org/10.1083/jcb.201805044. PMID: 31113824.

46. Wainman A, Buster DW, Duncan T, Metz J, Ma A, Sharp D, et al. A new Augmin subunit, Msd1, demonstrates the importance of mitotic spindle-templated microtubule nucleation in the absence of functioning centrosomes. Genes Dev. 2009;23(16):1876–81. https://doi.org/10.1101/gad.532209. PMID: 19684111.

47. Akhmanova A, Steinmetz MO. Control of microtubule organization and dynamics: two ends in the limelight. Nat Rev Mol Cell Biol. 2015;16(12):711–26. https://doi.org/10.1038/nrm4084. PMID: 26562752.

48. Akhmanova A, Steinmetz MO. Microtubule minus-end regulation at a glance. J Cell Sci. 2019;132(11). https://doi.org/10.1242/jcs.227850. PMID: 31175152.

49. Ems-McClung SC, Walczak CE. Kinesin-13s in mitosis: Key players in the spatial and temporal organization of spindle microtubules. Semin Cell Dev Biol. 2010;21(3):276–82. https://doi.org/10.1016/j.semcdb.2010.01.016. PMID: 20109574.

50. Goshima G, Wollman R, Stuurman N, Scholey JM, Vale RD. Length control of the metaphase spindle. Curr Biol. 2005;15(22):1979–88. https://doi.org/10.1016/j.cub.2005.09.054. PMID: 16303556.

51. Morales-Mulia S, Scholey JM. Spindle pole organization in Drosophila S2 cells by dynein, abnormal spindle protein (Asp), and KLP10A. Mol Biol Cell. 2005;16(7):3176–86. https://doi.org/10.1091/mbc.e04-12-1110. PMID: 15888542.

52. Buster DW, Zhang D, Sharp DJ. Poleward tubulin flux in spindles: regulation and function in mitotic cells. Mol Biol Cell. 2007;18(8):3094–104. https://doi.org/10.1091/mbc.e06-11-0994. PMID: 17553931.

53. Pavlova GA, Razuvaeva AV, Popova JV, Andreyeva EN, Yarinich LA, Lebedev MO, et al. The role of Patronin in Drosophila mitosis. BMC Mol Cell Biol. 2019;20(Suppl 1):7. https://doi.org/10.1186/s12860-019-0189-0. PMID: 31284878.

54. Goshima G, Vale RD. Cell cycle-dependent dynamics and regulation of mitotic kinesins in Drosophila S2 cells. Mol Biol Cell. 2005;16(8):3896–907. https://doi.org/10.1091/mbc.e05-02-0118. PMID: 15958489.

55. Rogers GC, Rogers SL, Schwimmer TA, Ems-McClung SC, Walczak CE, Vale RD, et al. Two mitotic kinesins cooperate to drive sister chromatid separation during anaphase. Nature. 2004;427(6972):364–70. https://doi.org/10.1038/nature02256. PMID: 14681690.

56. Laycock JE, Savoian MS, Glover DM. Antagonistic activities of Klp10A and Orbit regulate spindle length, bipolarity and function in vivo. J Cell Sci. 2006;119(Pt 11):2354–61. https://doi.org/10.1242/jcs.02957. PMID: 16723741.

57. Renda F, Pellacani C, Strunov A, Bucciarelli E, Naim V, Bosso G, et al. The Drosophila orthologue of the INT6 onco-protein regulates mitotic microtubule growth and kinetochore structure. PLoS Genet. 2017;13(5):e1006784. https://doi.org/10.1371/journal.pgen.1006784. PMID: 28505193.

58. Jiang K, Rezabkova L, Hua S, Liu Q, Capitani G, Altelaar AFM, et al. Microtubule minus-end regulation at spindle poles by an ASPM-katanin complex. Nat Cell Biol. 2017;19(5):480–92. https://doi.org/10.1038/ncb3511. PMID: 28436967.

59. Wakefield JG, Bonaccorsi S, Gatti M. The Drosophila protein Asp is involved in microtubule organization during spindle formation and cytokinesis. J Cell Biol. 2001;153(4):637–48. https://doi.org/10.1083/jcb.153.4.637. PMID: 11352927.

60. Ito A, Goshima G. Microcephaly protein Asp focuses the minus ends of spindle microtubules at the pole and within the spindle. J Cell Biol. 2015;211(5):999–1009. https://doi.org/10.1083/jcb.201507001. PMID: 26644514.

61. Maiato H, Khodjakov A, Rieder CL. Drosophila CLASP is required for the incorporation of microtubule subunits into fluxing kinetochore fibres. Nat Cell Biol. 2005;7(1):42–7. https://doi.org/10.1038/ncb1207. PMID: 15592460.

62. Inoue YH, do Carmo Avides M, Shiraki M, Deak P, Yamaguchi M, Nishimoto Y, et al. Orbit, a novel microtubule-associated protein essential for mitosis in Drosophila melanogaster. J Cell Biol. 2000;149(1):153–66. https://doi.org/10.1083/jcb.149.1.153. PMID: 10747094.

63. Lemos CL, Sampaio P, Maiato H, Costa M, Omel’yanchuk LV, Liberal V, et al. Mast, a conserved microtubule-associated protein required for bipolar mitotic spindle organization. EMBO J. 2000;19(14):3668–82. https://doi.org/10.1093/emboj/19.14.3668. PMID: 10899121.

64. Rogers SL, Rogers GC, Sharp DJ, Vale RD. Drosophila EB1 is important for proper assembly, dynamics, and positioning of the mitotic spindle. J Cell Biol. 2002;158(5):873–84. https://doi.org/10.1083/jcb.200202032. PMID: 12213835.

65. Goshima G, Wollman R, Goodwin SS, Zhang N, Scholey JM, Vale RD, et al. Genes required for mitotic spindle assembly in Drosophila S2 cells. Science. 2007;316(5823):417–21. https://doi.org/10.1126/science.1141314. PMID: 17412918.

66. Zhang G, Breuer M, Forster A, Egger-Adam D, Wodarz A. Mars, a Drosophila protein related to vertebrate HURP, is required for the attachment of centrosomes to the mitotic spindle during syncytial nuclear divisions. J Cell Sci. 2009;122(Pt 4):535–45. https://doi.org/10.1242/jcs.040352. PMID: 19174464.

67. Goshima G. Identification of a TPX2-like microtubule-associated protein in Drosophila. PLoS One. 2011;6(11):e28120. https://doi.org/10.1371/journal.pone.0028120. PMID: 22140519.

68. Pavlova GA, Galimova YA, Popova YV, Munzarova AF, Razuvaeva AV, Alekseeva AL, et al. Factors Governing the Pattern of Spindle Microtubule Regrowth after Tubulin Depolymerization. Tsitologiia. 2016;58(4):299–303. PMID: 30191704.

69. Goshima G, Vale RD. The roles of microtubule-based motor proteins in mitosis: comprehensive RNAi analysis in the Drosophila S2 cell line. J Cell Biol. 2003;162(6):1003–16. https://doi.org/10.1083/jcb.200303022. PMID: 12975346.

70. Reis R, Feijao T, Gouveia S, Pereira AJ, Matos I, Sampaio P, et al. Dynein and mast/orbit/CLASP have antagonistic roles in regulating kinetochore-microtubule plus-end dynamics. J Cell Sci. 2009;122(Pt 14):2543–53. https://doi.org/10.1242/jcs.044818. PMID: 19549687.

71. Yang CP, Fan SS. Drosophila mars is required for organizing kinetochore microtubules during mitosis. Exp Cell Res. 2008;314(17):3209–20. https://doi.org/10.1016/j.yexcr.2008.08.004. PMID: 18761010.

72. Saunders RD, Avides MC, Howard T, Gonzalez C, Glover DM. The Drosophila gene abnormal spindle encodes a novel microtubule-associated protein that associates with the polar regions of the mitotic spindle. J Cell Biol. 1997;137(4):881–90. https://doi.org/10.1083/jcb.137.4.881. PMID: 9151690.

73. Schoborg T, Zajac AL, Fagerstrom CJ, Guillen RX, Rusan NM. An Asp-CaM complex is required for centrosome-pole cohesion and centrosome inheritance in neural stem cells. J Cell Biol. 2015;211(5):987–98. https://doi.org/10.1083/jcb.201509054. PMID: 26620907.

74. Cheeseman IM, MacLeod I, Yates JR, 3rd, Oegema K, Desai A. The CENP-F-like proteins HCP-1 and HCP-2 target CLASP to kinetochores to mediate chromosome segregation. Curr Biol. 2005;15(8):771–7. https://doi.org/10.1016/j.cub.2005.03.018. PMID: 15854912.

75. Maffini S, Maia AR, Manning AL, Maliga Z, Pereira AL, Junqueira M, et al. Motor-independent targeting of CLASPs to kinetochores by CENP-E promotes microtubule turnover and poleward flux. Curr Biol. 2009;19(18):1566–72. https://doi.org/10.1016/j.cub.2009.07.059. PMID: 19733075.

76. Hannak E, Heald R. Xorbit/CLASP links dynamic microtubules to chromosomes in the Xenopus meiotic spindle. J Cell Biol. 2006;172(1):19–25. https://doi.org/10.1083/jcb.200508180. PMID: 16390996.

77. Kitamura E, Tanaka K, Komoto S, Kitamura Y, Antony C, Tanaka TU. Kinetochores generate microtubules with distal plus ends: their roles and limited lifetime in mitosis. Dev Cell. 2010;18(2):248–59. https://doi.org/10.1016/j.devcel.2009.12.018. PMID: 20159595.

78. Buffin E, Emre D, Karess RE. Flies without a spindle checkpoint. Nat Cell Biol. 2007;9(5):565–72. https://doi.org/10.1038/ncb1570. PMID: 17417628.

79. Maiato H, Fairley EA, Rieder CL, Swedlow JR, Sunkel CE, Earnshaw WC. Human CLASP1 is an outer kinetochore component that regulates spindle microtubule dynamics. Cell. 2003;113(7):891–904. https://doi.org/10.1016/s0092-8674(03)00465-3. PMID: 12837247.

80. Liu P, Wurtz M, Zupa E, Pfeffer S, Schiebel E. Microtubule nucleation: The waltz between gamma-tubulin ring complex and associated proteins. Curr Opin Cell Biol. 2021;68:124–31. https://doi.org/10.1016/j.ceb.2020.10.004. PMID: 33190097.

81. Sillje HH, Nagel S, Korner R, Nigg EA. HURP is a Ran-importin beta-regulated protein that stabilizes kinetochore microtubules in the vicinity of chromosomes. Curr Biol. 2006;16(8):731–42. https://doi.org/10.1016/j.cub.2006.02.070. PMID: 16631580.

82. Jiang K, Faltova L, Hua S, Capitani G, Prota AE, Landgraf C, et al. Structural Basis of Formation of the Microtubule Minus-End-Regulating CAMSAP-Katanin Complex. Structure. 2018;26(3):375–82 e4. https://doi.org/10.1016/j.str.2017.12.017. PMID: 29395789.

83. Pavlova GA, Popova JV, Andreyeva EN, Yarinich LA, Lebedev MO, Razuvaeva AV, et al. RNAi-mediated depletion of the NSL complex subunits leads to abnormal chromosome segregation and defective centrosome duplication in Drosophila mitosis. PLoS Genet. 2019;15(9):e1008371. https://doi.org/10.1371/journal.pgen.1008371. PMID: 31527906.

84. Bustin SA, Benes V, Garson JA, Hellemans J, Huggett J, Kubista M, et al. The MIQE guidelines: minimum information for publication of quantitative real-time PCR experiments. Clin Chem. 2009;55(4):611–22. https://doi.org/10.1373/clinchem.2008.112797. PMID: 19246619.

85. Yang X, Mao F, Lv X, Zhang Z, Fu L, Lu Y, et al. Drosophila Vps36 regulates Smo trafficking in Hedgehog signaling. J Cell Sci. 2013;126(Pt 18):4230–8. https://doi.org/10.1242/jcs.128603. PMID: 23843610.

86. Mackay TF, Richards S, Stone EA, Barbadilla A, Ayroles JF, Zhu D, et al. The Drosophila melanogaster Genetic Reference Panel. Nature. 2012;482(7384):173–8. https://doi.org/10.1038/nature10811. PMID: 22318601.

87. Huang W, Massouras A, Inoue Y, Peiffer J, Ramia M, Tarone AM, et al. Natural variation in genome architecture among 205 Drosophila melanogaster Genetic Reference Panel lines. Genome Res. 2014;24(7):1193–208. https://doi.org/10.1101/gr.171546.113. PMID: 24714809.

88. Somma MP, Fasulo B, Cenci G, Cundari E, Gatti M. Molecular dissection of cytokinesis by RNA interference in Drosophila cultured cells. Mol Biol Cell. 2002;13(7):2448–60. https://doi.org/10.1091/mbc.01-12-0589. PMID: 12134082.

89. Spitzer M, Wildenhain J, Rappsilber J, Tyers M. BoxPlotR: a web tool for generation of box plots. Nat Methods. 2014;11(2):121–2. https://doi.org/10.1038/nmeth.2811. PMID: 24481215.

90. Schneider CA, Rasband WS, Eliceiri KW. NIH Image to ImageJ: 25 years of image analysis. Nat Methods. 2012;9(7):671–5. https://doi.org/10.1038/nmeth.2089. PMID: 22930834.

