## Supplementary material for "Genetic control of kinetochore-driven microtubule growth in *Drosophila* mitosis": S1 Table

| Targeted gene | Relative mRNA level after RNAi  %  (± SEM) | Centrosomes/spindle poles | | | Distribution of cells by stages of  mitosis (%) | | | |
| --- | --- | --- | --- | --- | --- | --- | --- | --- |
|  |  | # of cells scored  (1) | Monopolar  % | >2 centros.  % | # of cells scored (2) | Prometa and Meta | Ana and telo | PMLES |
| None (control) | - - - | 616 | 3.3 | 7.6 | 549 | 59.4 | 35.0 | 5.6 |
| *asp* | 6.4 (± 0.9) | 262 | 4.2 | 5.0 | 238 | 69.6* | 19.0* | 11.4* |
| *Dgt6* | 4.8 (±1.0) | 340 | 14.4* | 4.7 | 275 | 74.8* | 8.2* | 17.0* |
| *Eb1* | 2.8 (± 0.5) | 356 | 3.1 | 10.4 | 308 | 62.9 | 25.8* | 11.3* |
| *Klp10A* | 8.0 (± 1.3) | 356 | 16.9* | 9.8 | 261 | 62.7 | 27.7§ | 9.6§ |
| *mars* | 3.0 (± 1.0) | 338 | 6.8§ | 7.4 | 290 | 67.2§ | 24.5* | 8.3 |
| *mast* | 8.4 (± 2.5) | 425 | 49.6* | 8.0 | 181 | 84.5* | 7.7* | 7.8 |
| *mei-38* | 5.0 (± 1.4) | 458 | 23.8* | 15.7* | 277 | 70.1* | 21.2* | 8.7 |
| *Patronin* | 28.0 (± 3.0) | 261 | 6.9§ | 15.7* | 202 | 58.6 | 34.8 | 6.6 |

**Table S1. RNAi efficiency and frequencies of mitotic figures observed after RNAi against *asp*, *dgt6*, *Eb1, Klp10A, mars, mast*, *mei-38* and *Patronin***. The analysis of the mRNA levels after RNAi was performed by RT-qPCR and reported as the median value across the replicates ± SEM. The frequencies of cells with monopolar spindles and multiple (> 2) centrosomes were determined by examining all mitotic cells (# of cells scored, 1). We did not try to distinguish between monopolar spindles with a single centrosome and monopolars with 2 centrosomes at center of a monaster. The frequencies of the different mitotic figures were determined by examining only bipolar spindles with two centrosomes (# of cells scored, 2). Prometa, prometaphases; Meta, metaphases; Ana, anaphases; Telo, telophases; PMLES, prometaphase-like cells with elongated spindles. § and *, significant in χ^2^ test with p ≤ 0.05, and ≤ 0.01, respectively.
