## Supplementary material for "Genetic control of kinetochore-driven microtubule growth in *Drosophila* mitosis": S2 Table

**Table S2.** **dsRNAs used for RNA interference.**

| **dsRNA probe** | | | **Sequences of primers (5′->3′) used to amplify**  **DNA templates for dsRNA synthesis (*)** |
| --- | --- | --- | --- |
| **Target gene** | **Length,**  **bp** | **Reference** |  |
| *asp* | 959 | [1] | CTGCGATCTTTCTTCAG  AGATGATTACGCCAATGC |
| *dgt6* | 785 | [2] | ATCGGACCATAA  TTGTTCTCGGCT |
| *Eb1* | 322 | This study | TGCTGCCGCGCA  TGGTGCATCGTCAGGC |
| *Klp10A* | 618 | This study | TTGCTGTCCATC  CGATCCTTGTCT |
| *mars* | 576 | [2] | TGAACTCGCGCT  AACTGCTGCAGA |
| *mast* | 913 | This study | CGTTCTCGGAGC  ATGTCCACGAAG |
| *mei-38* | 614 | [2] | CGTCCAAGGATG  TGGAAGGGTCTG |
| *Patronin* | 923 | [1] | CGAGCTACAGCACCTGTTTC  TGACTGATTGCTGACATCGTCC |

(*) Each primer contained the following additional sequence of the T7 RNA polymerase-binding site at the 5′ end (5′->3′): TAATACGACTCACTATAGGGAGG.

**References**

1.	Pavlova GA, Razuvaeva AV, Popova JV, Andreyeva EN, Yarinich LA, Lebedev MO, et al. The role of Patronin in Drosophila mitosis. BMC Mol Cell Biol. 2019;20(Suppl 1):7. https://doi.org/10.1186/s12860-019-0189-0. PMID: 31284878.

2.	Somma MP, Ceprani F, Bucciarelli E, Naim V, De Arcangelis V, Piergentili R, et al. Identification of Drosophila mitotic genes by combining co-expression analysis and RNA interference. PLoS Genet. 2008;4(7):e1000126. https://doi.org/10.1371/journal.pgen.1000126. PMID: 18797514.
