## Supplementary material for "Genetic control of kinetochore-driven microtubule growth in *Drosophila* mitosis": S3 Table

**Table S3. Primers used for assessment of the RNAi-mediated gene silencing by RT-qPCR**

| **Target gene** | **Primer sequence (5′->3′)** | **Primers efficiency,**  **%** | **Amplicon size (from cDNA), bp** |
| --- | --- | --- | --- |
| *asp* | AAGTCGATTGGATCGTCTTTC | 105.1 | 155 |
|  | AATTTAGGATGATCCGGCTG |  |  |
| *dgt6* | AACAGCTTACTCGCACCTGC | 101.3 | 118 |
|  | GCATGGGATCGTTGATCTTG |  |  |
| *Eb1* | TCTGCACAGGTGCAGCTTACTGTCA | 102.9 | 140 |
|  | CTTCTTGAAGCCCGCCTGCA |  |  |
| *Klp10A* | GCTGAGCGAACACGAGATGT | 100.2 | 115 |
|  | CAGTGTGGCATTAACGGTGC |  |  |
| *mars* | CAGACGACGGTTAAAGAAGACA | 105.9 | 121 |
|  | GCGAGACACTAACAAACGGA |  |  |
| *mast* | AGACGCTGAGAAATAAACTAGATGC | 106.6 | 147 |
|  | GCAGCTTTGGTGCATGTGTA |  |  |
| *mei-38* | TTTGGCCAGTCGGACTTCGA | 101.7 | 165, 168 |
|  | CTGTTCCCGGCCGATCTTCT |  |  |
| *Patronin* | TTTTCAAATACAACTCAGGAGGCA | 99.0 | 82 |
|  | ATTGTGAAGGCGTCGATGGT |  |  |
| *RpL32*  (as reference) | CTAAGCTGTCGCACAAATGG | 100.0 | 148 |
|  | AGGAACTTCTTGAATCCGGTG |  |  |
